# Stimulation of Neurons and Astrocytes via Temporally Interfering Electric Fields

**DOI:** 10.1101/2023.10.30.564774

**Authors:** Annika Ahtiainen, Lilly Leydolph, Jarno M.A. Tanskanen, Alexander Hunold, Jens Haueisen, Jari A.K. Hyttinen

## Abstract

A novel non-invasive electrical stimulation method, temporal interference stimulation (TIS), has been proposed to enable spatially steerable stimulation and to selectively activate different brain regions. However, TIS has not been studied with cell cultures that can provide an indepth assessment of its effects on neuronal cells. We successfully established an *in vitro* TIS setup using rat cortical neurons on microelectrode arrays. Stimulations involved 1) TIS with 653 Hz and 643 Hz, resulting in a 10 Hz frequency envelope, 2) low-frequency stimulation (LFS) at 10 Hz, 3) high-frequency stimulation (HFS) at 653 Hz, and 4) no electrical stimulation (control/sham). HFS and LFS had the least effect on neuronal activity, but TIS elicited neuronal electrophysiological responses, especially 24 hours after stimulation. We also assessed TIS in neuron-astrocyte co-cultures. Interestingly, co-cultures seemed to counteract the TIS effects. Hence, our findings deepen the current understanding of the electrical modulation of neurons during TIS.

**Highlights:**

- Temporal interference stimulation (TIS) is a new technique for brain stimulation.
- TIS setup was established and combined with microelectrode array (MEA) system.
- Rat cortical neurons and astrocytes were stimulated with TIS *in vitro*.
- Neurons responded to TIS more prominently without astrocyte support.

## Introduction

A promising new technique known as temporal interference stimulation (TIS) has been proposed to open new possibilities in the treatment of neurological conditions. TIS is a non-invasive electrical stimulation (ES) method that promises targeting of neurons at specific depths (Grossman et al., 2017). Traditional non-invasive brain stimulation methods, such as transcranial magnetic stimulation (TMS) and transcranial electrical current stimulation (tES), have lacked selectivity and effectiveness to therapeutically address target structures at necessary depth. Hence, the primary option has been invasive deep brain stimulation (DBS), despite its associated risks (Krauss et al., 2021; Lozano et al., 2019; Videnovic and Metman, 2008). TIS, however, possesses considerable promise, as it can be steered to affect only specific deep regions without modulating the superficial layers of the brain (Dmochowski and Bikson, 2017; Grossman et al., 2017). TIS has demonstrated the ability to target deep brain structures and modulate brain functions *in vivo* (Violante et al., 2023). Specifically, TIS enhances motor learning and memory performance in humans by non-invasively modulating memory activity in the striatum and hippocampus (Violante et al., 2023; Wessel et al., 2023).

TIS is established using two high-frequency sinusoidal currents with different frequencies (i.e., f_2_ > f_1_> 600 Hz) that, when interfering in the volume conductor, form a signal with a low-frequency “envelope” Δf (Δf=f_2_-f_1_), usually between 1 and 100 Hz. As the high-frequency fields are above the frequency range defined by the cell membrane time constant, cells – and neurons – do not react individually to the high frequencies. However, the low-frequency interference field of the resulting beat frequency Δf has the potential to influence neuronal transmembrane potential (Bikson et al., 2004; Deans et al., 2007; Karimi et al., 2019). Therefore, TIS has recently drawn attention from researchers because it could offer new ways of modulating the neuronal systems and treating neurological conditions such as Parkinson’s disease and chronic pain (Grossman et al., 2018; Guo et al., 2023; Mirzakhalili et al., 2020; Opitz and Tyler, 2017; Rampersad et al., 2019; Song et al., 2021; Zhu and Yin, 2023). Thus far, TIS has been demonstrated using computational studies and *in vivo* applications with rodents (Carmona-Barrón et al., 2023; Grossman et al., 2017) and human subjects (Ma et al., 2022; Violante et al., 2023; Zhu et al., 2022). However, it has not been tested with *in vitro* cell cultures, where the effects of electrical field interference (EFI) on neuronal cells and their interactions could be assessed more in depth. Furthermore, the role of astrocytes during TIS remains unexplored; astrocytes are the “silent influencers” in the brain that have lately drawn great interest due to their pivotal role in neuronal signaling, homeostasis, and diseases.

In this study, we introduce an *in vitro* TIS platform for neuronal cell cultures conducted similarly to that of Grossman et al. for brain stimulation (Grossman et al., 2017). The stimulation setup combined an EFI current stimulator, electrodes for the stimulation, and microelectrode array (MEA) technology, capable of recording and monitoring the resulting stimulus fields and neuronal electrical activity in real time (Thomas et al., 1972). We utilized different ES paradigms to compare the effects of various stimulation modalities on neural cultures *in vitro*. We have previously used similar defined neuron and neuron-astrocyte co-cultures to study the effects of pharmacological stimulation (Ahtiainen et al., 2023, 2021). Neurons and astrocytes are the two major cell types in the brain. While neurons transmit electrical signals, astrocytes regulate neuronal activity by clearing and secreting neurotransmitters, buffering ions, and hence directly affecting neuronal electrical activity and brain homeostasis (Bellot-Saez et al., 2017; McNeill et al., 2021). Astrocytes have been implicated in multiple neurogenerative diseases, such as Alzheimer’s and Parkinson’s disease, the symptoms of which could be alleviated with TIS (Brandebura et al., 2023; Ramos-Gonzalez et al., 2021; Wang and Ye, 2021).

Our aim was to establish EFI stimulation *in vitro* and investigate how neurons respond to TIS and its low-frequency (LFS) and high-frequency (HFS) counterparts, both in the short term and over an extended period. We observed that TIS highly affected the electrophysiological activity of neurons, especially 24 hours after stimulation, while HFS and LFS had very different effects. Interestingly, neurons with comprehensive astrocytic support responded to TIS differently than neuronal cultures with less astrocytic impact. Here, we demonstrate the usability of the *in vitro* EFI and show that it can be applied and controlled to study its effects on neural cell cultures. Moreover, we measured the stimulus electric fields and demonstrated the spatial steerability of our TIS setup. Our study paves the way for in evaluating the effects of TIS on neuronal cells and diseases, opens the way to assess its potential therapeutic applications and sheds light on the role of astrocytes in modulating neuronal responses to it.

## Results

### Electrical stimulation concept and validation on microelectrode arrays

We established a microelectrode array (MEA) stimulation and recording setup by combining a neuroConn current stimulator with a Multi Channel Systems (MCS) MEA2100 system that can record neuronal electrical activity. The stimulus was conducted to cells located at the center of the MEA electrode area via platinum rod electrodes that were submerged into the cell medium through a 3D-printed cap (**Supplementary Figure S1A-C)**. An illustration of the stimulation and recording setup is shown in **Figure 1A**. Before the cell experiments, the system and applied stimulation currents were validated with measurements using an oscilloscope (**Supplementary Figure S1D-E**).

**Figure 1.**
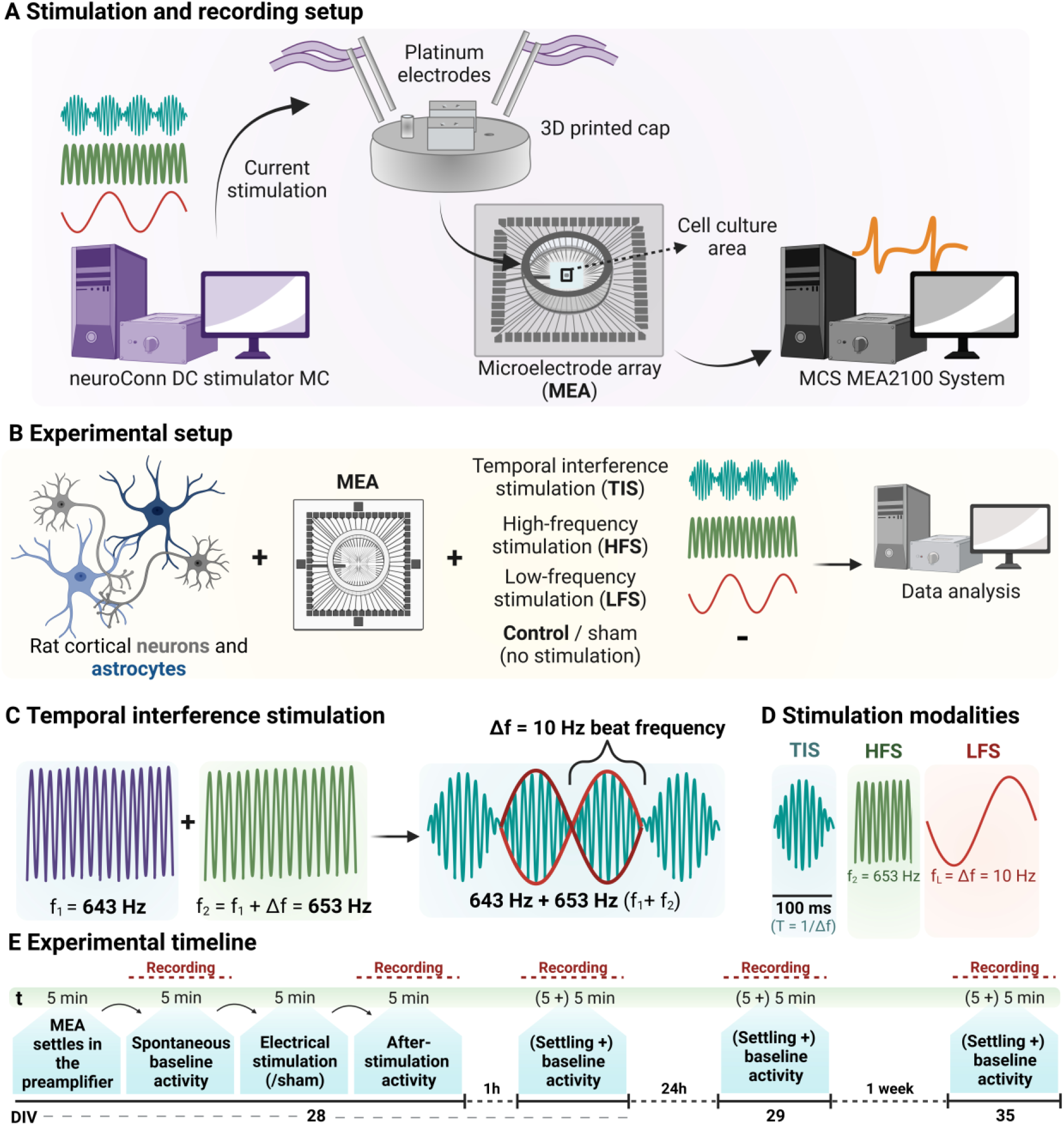
Temporal interference stimulation and the experimental timelines. (**A**) Neuron and neuron-astrocyte co-cultures were stimulated using a neuroConn stimulator. The stimulus was conducted via platinum rod electrodes submerged in the cell medium in the microelectrode arrays (MEAs). The data were collected using a MEA2100 system from Multi Channel Systems (MCS) GmbH. (**B**) Rat cortical neurons and astrocytes are plated on MEAs. Neuronal cultures were stimulated with TIS, LFS, HFS, and control, and the neuron-astrocyte co-cultures (∼1:1) were stimulated with TIS and control/sham. (**C**) The concept of temporal interference stimulation TIS is formed by two different high-frequency signals (f_1_ and f_2_) that result in a beat frequency Δf with stimulation envelopes corresponding to the LFS signal (10 Hz). (**D**) Different stimulation modalities (TIS, HFS, and LFS) were used in the experiment. Whereas the LFS consisted of one cycle of stimulation at 10 Hz, the TIS consisted of high frequencies within the formed 10 Hz envelope, as in the HFS. (Representative signal waveforms within TIS and HFS signal cycles are not shown to scale.) (**E**) At 28 days in vitro (DIV), cultures were recorded after five-minute settling and after electrical stimulation (or sham). The activity was recorded immediately after stimulation, and one hour, one day, and one week after stimulation. The figure was created with Biorender.com.

The stimulation and recording setups were used to stimulate rat cortical neurons and astrocytes on MEAs (**Figure 1B**). We used four different stimulation groups – temporal interference stimulation (TIS), high-frequency stimulation (HFS), low-frequency stimulation (LFS), and control/sham for neurons (n=16 MEAs). Neuron-astrocyte co-cultures were stimulated with TIS or control/sham (n=8 MEAs). TIS is composed of two slightly different high-frequency currents (f_1_ and f_2_) that produce a low beat frequency (Δf), which falls into the range to which neurons can respond (**Figure 1C**). To compare the effects of TIS, HFS and LFS with frequencies corresponding to that of TIS were also used to stimulate neurons (**Figure 1D**). The electrical stimulus was applied for five minutes at 28 days *in vitro* (DIV). Neuronal electrical activity was recorded before stimulation/sham, immediately after, one hour after, 24 hours after, and one week after stimulation/sham (**Figure 1E**).

To achieve the stimulation setup for electrical stimulation (ES), a two-channel electrical current stimulation system was established. **Figure 2A** shows the schematics of the TIS stimulation setup with two electrode pairs fed with currents of 450 µA. The stimulation reached the MEA electrode area, where the cells were plated, from two sides through placement of the platinum rod electrode pairs at a 25° angle (**Figure 2B**). Prior to the stimulation of the cells, the stimulation system that combined the neuroConn DC-STIMULATOR MC and the MEA2100 system was evaluated with MEAs without any cell cultures. Using only the cell culture medium, all stimulations (TIS, HFS, and LFS) were also observed in the MEA2100 recordings (**Figure 2C**). **Figure 2D** shows a 1-second close-up of each of the stimulations. The LFS had the lowest frequency at 10 Hz, the HFS had high-frequency characteristics at 653 Hz, and the TIS had 10 Hz envelopes, along with high-frequency signals within the envelope, as expected.

**Figure 2.**
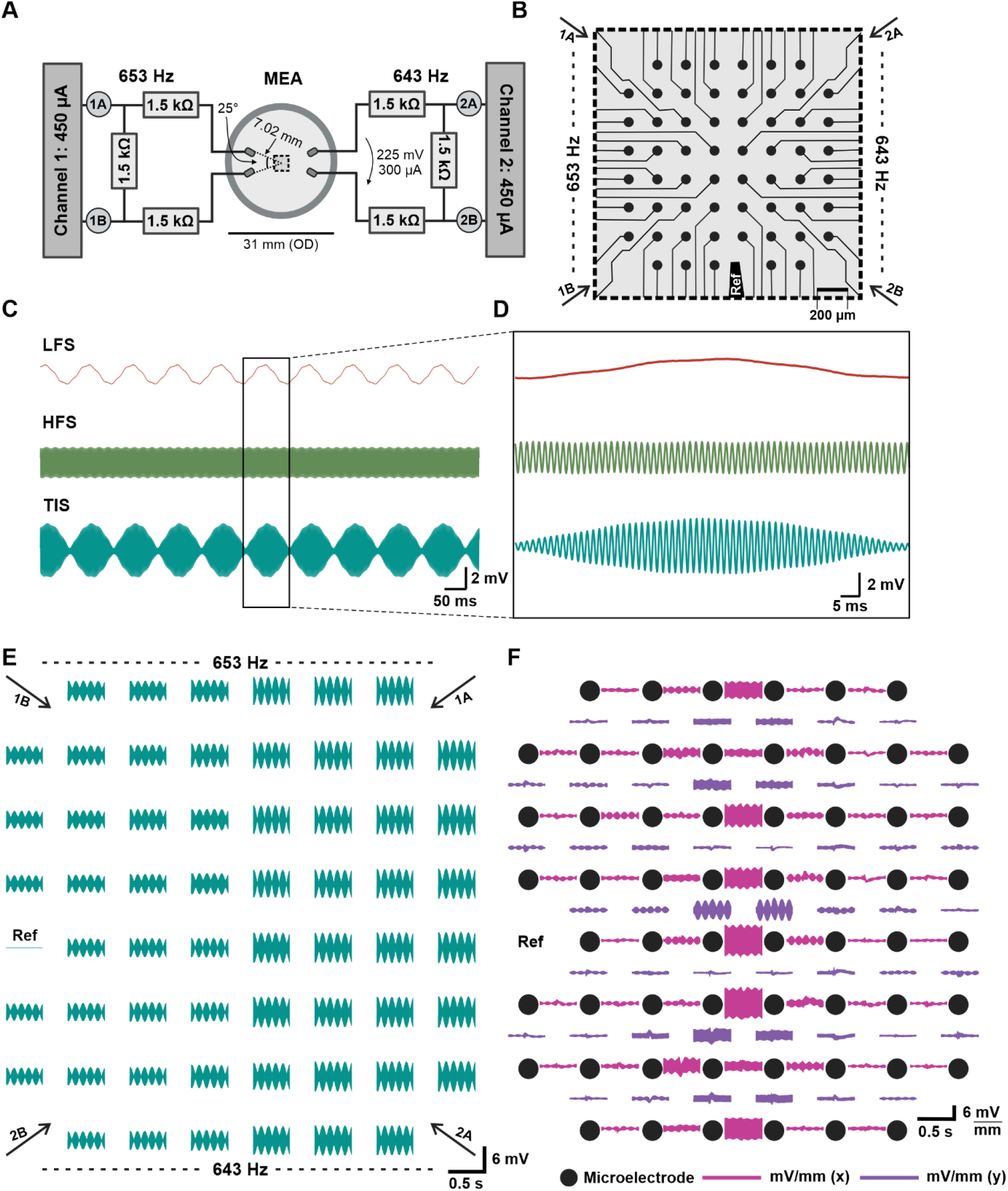
Electrical stimulation system and its validation. (**A**) Schematic of the experimental setup, including V_ch1_=V_ch2_=450 µA with 1.5 kΩ resistors. A microelectrode array was located in between the two channels, and the stimulus reached the MEA at a 25° angle. (**B**) MEA electrode area. Microelectrodes are depicted in dark gray circles, and the reference electrode is indicated as a black bar. The distance between each microelectrode is 200 µm, and the electrode diameter is 30 µm. The MEA electrode area in **A,B** is at a 90° angle with the electrode area shown in **E,F**. (**C**) 1-second recordings of the low-frequency stimulation (LFS), high-frequency stimulation (HFS), and temporal interference stimulation (TIS) on the MEAs. (**D**) 100 ms close-ups from each of the stimulations (HFS/LFS/TIS). (**E**) A representative MEA2100 recording of the whole electrode area of a TIS-stimulated co-culture on the MEA. Arrows indicate the direction of the applied stimulus and the applied frequencies. (**F**) The electric field potentials (indicated with pink and purple traces) between two adjacent electrodes in both horizontal and vertical directions were lower at the sides of the electrode area and highest in the middle of it, suggesting the steerability of our TIS system. The microelectrodes are depicted as dark gray circles.

Next, we evaluated the system with cell cultures; **Figure 2E** shows the entire electrode area of one representative TI-stimulated co-culture on the MEA. Depending on the electrode in question, the TIS amplitudes ranged from -2.1±0.4 to 2.1±0.4 mV, with a peak-to-peak amplitude of 4.2 mV (**Figure 2E, Table 1**). The TIS amplitude was the lowest around the reference electrode (electrode number 15), which was internally grounded in the system. The amplitudes were the highest further away from the reference electrode. We also calculated the differences in the local electric potentials between two adjacent electrodes in both horizontal and vertical directions (mV/mm [x] and mV/mm [y], respectively) for the same TIS-stimulated co-culture MEA, shown in **Figure 2E**. The electric potential differences sensed by the cells also varied within the electrode area, being the highest in the middle of the electrode area in both vertical and horizontal directions and lowest at the edges of the electrode area (**Figure 2F**). This indicates that the TIS steers the stimulation to a specific target area in our *in vitro* system. The representative signal measurements and field strengths for all stimulation modalities are visible in **Supplementary Figure S3**.

**Table 1.** Minimum and maximum amplitude values (±standard deviation [SD]) of TIS and the electric potential differences (mV/mm) across electrodes for a representative co-culture MEA during TIS.

|  |  | TIS (mV) | mV/mm (x) | mV/mm (y) |
| --- | --- | --- | --- | --- |
| Minimum | Mean±SD | $-2.1 \pm 0.4$ | $-1.8 \pm 1.2$ | $-1.4 \pm 0.7$ |
|  | Median | -2.3 | -1.4 | -1.2 |
| Maximum | Mean±SD | $2.1 \pm 0.4$ | $1.8 \pm 1.2$ | $1.4 \pm 0.8$ |
|  | Median | 2.2 | 1.3 | 1.2 |

### Maturation of the neural cultures

Neuron and neuron-astrocyte co-cultures were followed for 35 days *in vitro* (DIV) on MEAs (representative images of the cultures at DIV28 in **Figure 3A**). In co-cultures, neurons and astrocytes were cultured in a ratio of ∼1:1, and the astrocytes were treated with ara-c (cytosine β-D-arabinofuranoside) to prevent their proliferation and to keep the cell numbers constant throughout the culture period. For both cultures, the number of neurons was the same, and only the proportion of astrocytes differed. **Figure 3B-C** shows the representative signal traces of both neurons and co-cultures at DIV28, with neurons exhibiting higher electrical activity.

**Figure 3.**
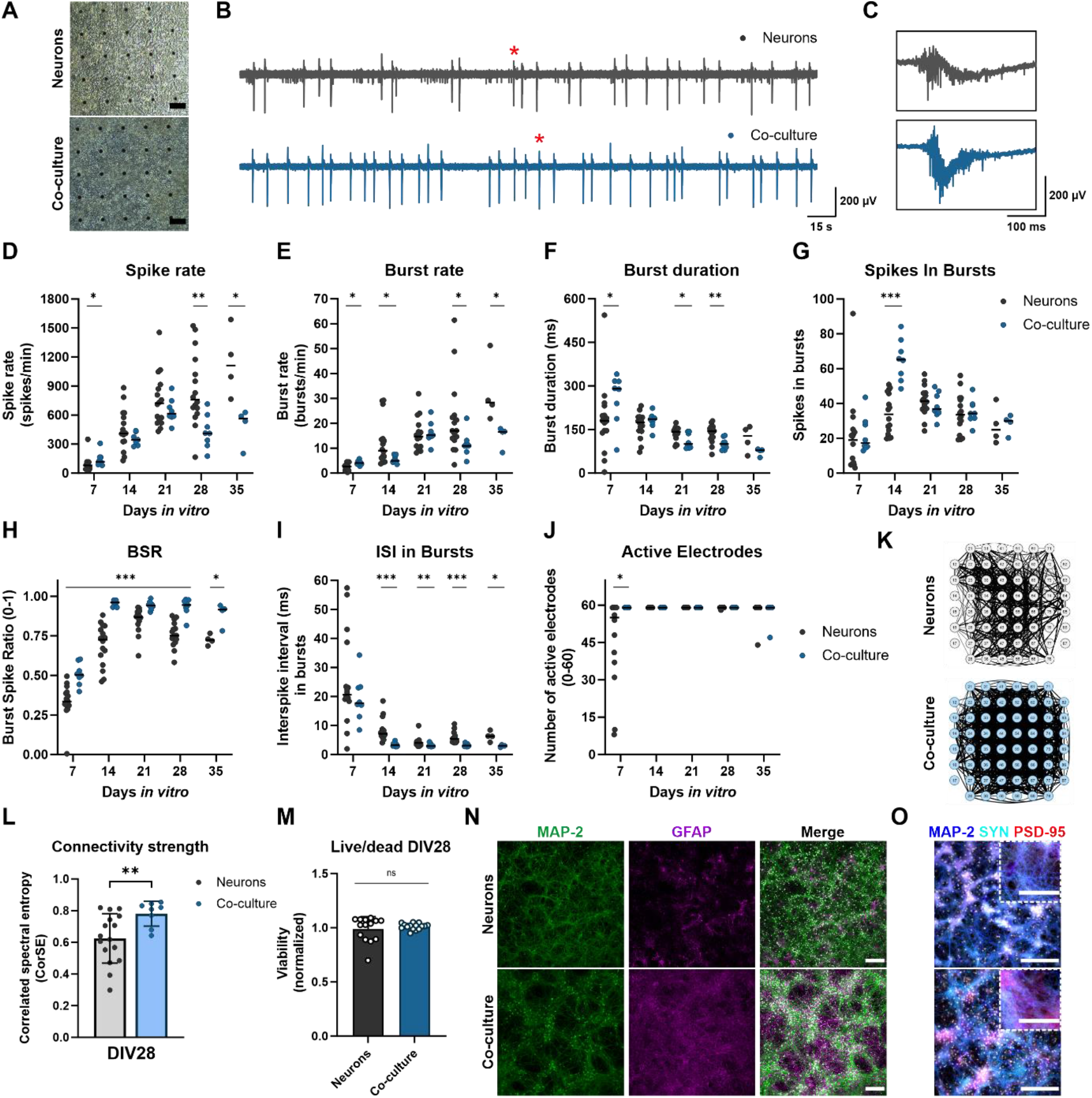
Characterization of the cultures. (**A**) Brightfield images of neuron- and neuron-astrocyte co-cultures on MEAs at DIV28. The scale bar is 100 µm. The black dots are the microelectrodes on the MEA surface. (**B-C**) Representative signal traces of neurons and co-cultures at DIV28. Red stars indicate the spots of the 300 ms closeups in **C**. (**D-J**) Electrophysiological characteristics of the cultures during five weeks in vitro. Each data point is represented, and the median is indicated with a black line. In general, neurons had higher electrical activity than co-cultures during the last weeks of culture but a lower burst spike ratio (BSR%) than co-cultures. (**K-L**) Correlation spectral entropy (CorSE) analysis revealed that the co-cultures had 25% greater functional connectivity than neurons at DIV28. (**M**) Live/dead assay of neuron and neuron-astrocyte co-cultures at DIV28. There were no differences in the viability of the cultures. Bar graphs in (**L-M**) show means±SD with data points. (**N**) Immunocytochemical (ICC) images of neurons (MAP2, microtubule associated protein 2) and astrocytes (GFAP, glial fibrillary acidic protein) at DIV29. Co-cultures had substantially more astrocytes than neuronal cultures without added astrocytes. Nucleus staining DAPI (4’,6-diamidino-2-phenylindole) is indicated in white. The scale bar is 200 µm. (**O**) ICC images of neurons (MAP2) and pre- and postsynaptic proteins (SYN, synaptophysin; PSD-95, respectively) at DIV29. Nucleus staining DAPI is indicated in white. The scale bar is 200 µm, and 100 µm for the close-ups. ^*^ p<0.05, ^**^p<0.01, ^***^p<0.001, ns=not significant. (n=16 neurons, n=8 co-cultures for MEAs, except for DIV35 n=4 MEAs for neurons and co-cultures. For live/dead, n=14/15 regions of interest [ROIs] in total.) All statistical tests were done using the Mann-Whitney U test.

The electrical activity of the cultures was evaluated weekly, starting with DIV7. The assessed electrophysiological features and their descriptions are listed in **Table 2**. All statistical tests and details of the maturation of the cultures can be found in **Supplementary Table S1**. At DIV7, co-cultures had higher activity than neurons (*p*=0.0192; Mann-Whitney U test). However, neurons had more spiking at DIV28 (*p*=0.0022) and DIV35 (*p*=0.0286) compared to co-cultures. For neuronal cultures, the spike rates and burst rates rose steadily until DIV35, whereas the co-cultures had their highest activity around DIV21 (**Figure 3D-E**). At DIV7, co-cultures had higher burst rates than neurons (*p*=0.0192), but after the first week *in vitro*, the neuronal cultures without astrocytes had higher burst rates than co-cultures (*p*=0.0382, *p*=0.0274, *p*=0.0286 at DIV14, 28 and 35, respectively).

**Table 2.** The assessed electrophysiological features of the cultures and their descriptions.

| Electrophysiological features | Description | Unit/scale |
| --- | --- | --- |
| **Number of active electrodes** | The number of active electrodes in MEA. An electrode is considered active if it has $\geq 10$ spikes/min (otherwise excluded from the analysis). | 0–60 |
| **Spike rate** | Number of spikes per minute (spike detection threshold $\pm 5\sigma$). | 1/min |
| **Burst rate** | Number of bursts per minute. Burst consists of $\geq 3$ spikes. | 1/min |
| **Burst duration** | The average duration of a burst. | ms |
| **Spikes in bursts** | The average number of spikes in bursts. | $\geq 0$ |
| **Burst spike ratio (BSR%)** | Percentage of spikes involved in bursts. | 0–1<br>(1=100%) |
| **Interspike interval (ISI) in bursts** | Average time between sequential spikes in bursts. | ms |
| **Correlated spectral entropy (CorSE)** | Evaluates MEA signal synchronicity by calculating the correlations of time-varying spectral entropies between the MEA channels (connection threshold $\geq 0.75$). | 0–1 |

For both neurons and co-cultures, the burst duration shortened toward DIV35 (**Figure 3F**), and co-cultures had shorter bursts than neurons, except at DIV7 (*p*=0.0274, *p*=0.0230, *p*=0.0071 for DIV7, DIV21, and DIV28, respectively). Also, the number of spikes in bursts reduced toward DIV35 and was significantly different between the cultures only at DIV14 (**Figure 3G**; *p*<0.001). The ratio of spikes involved in bursts was lower at DIV7 but rose rapidly already at DIV14 and was significantly different between the cultures every week (**Figure 3H**; *p*<0.001 for all except *p*=0.0286 at DIV35). Interspike interval (ISI) in bursts was lower for the co-cultures during the whole following period except for DIV7 (**Figure 3I**; *p*<0.001 at DIV14 and DIV28, and *p*=0.0057 and *p*=0.0285 at DIV21 and DIV35, respectively). The number of active electrodes was, except for one MEA, still 59 (out of 60) at DIV35 for both cultures and was different only for the cultures at DIV7 (**Figure 3J**; *p*=0.0158). Furthermore, differences in the connectivity of the neuron and neuron-astrocyte co-cultures were evaluated with correlation spectral entropy (CorSE) analysis. In general, the co-cultures had a 25% higher degree of functional connectivity than neurons at DIV28, indicating higher neuronal network connectivity for co-cultures across the MEA area (**Figure 3K-L**, *p*=0.0057; Mann-Whitney U test).

The viability of both cultures was also analyzed at DIV28. There was no significant difference in the viability of the cultures, and both cultures had very few dead cells (viability% for neurons 83±10% and 85±2% for co-cultures; **Figure 3M, Supplementary Figure S2A**). Furthermore, at DIV29, cultures were stained with immunocytochemistry (ICC), and the co-cultures had many more astrocytes than the neuronal cultures without separately added astrocytes, as expected (**Figure 3N**). Moreover, pre- and postsynaptic proteins, indicated by synaptophysin (SYN) and postsynaptic density protein-95 (PSD-95), were present in both cultures, implying functional connections between the neurites in both culture types (**Figure 3O**).

### Electrical stimulation of neurons *in vitro*

At DIV28, neurons were subjected to ES or sham for five minutes (n=4 MEAs/condition). The effects of ES on electrophysiology were evaluated by recording the activity for five minutes immediately, one hour, 24 hours, and one week after ES. To reliably assess the effects of TIS on cultures, neurons were also exposed to HFS and LFS stimulation with TIS characteristics. TIS comprised high-frequency (653+643 Hz) signals with a 10 Hz beat frequency that corresponded to the frequencies used for HFS and LFS stimulations (653 Hz and 10 Hz, respectively). Regarding the results, all statistical tests and details can be found in **Supplementary Tables S2-4**.

Prior to electrical stimulation, all the cultures had very similar activity except for HFS cultures, which had slightly elevated spike rates (+34%, *p*=0.0286; Mann-Whitney U test) and smaller BSRs than control (-8%, *p*=0.0286) (**Figure 4A,E**). Directly after the five-minute stimulation, LFS cultures experienced decreased spike rates (-26%), burst rates (-52%), elevated burst duration (+105%), number of spikes in bursts (+61%), and ISI (+41%) compared to control cultures (**Figure 4A-D,F**; *p*=0.0286 for all). As expected, HFS did not affect the neurons as much as the LFS, but the HFS cultures had decreased burst rates (-59%) and BSR (-20%) compared to control cultures right after the stimulation (**Figure 4B,E**; *p*=0.0286). Furthermore, one hour after stimulation, LFS and HFS showed decreased spiking (-51% and -37%, respectively), and bursting (-70% and -80%, respectively), increased burst duration (+227 for LFS and 171% for HFS) and BSRs (+22% for LFS and +17% for HFS), and an increased number of spikes in bursts (+178%) for the LFS (*p*=0.0286 for all). Moreover, TIS cultures had an elevated number of spikes in bursts (+159%) and BSR (+22%) one hour after the stimulation compared to the control cultures (*p*=0.0286), and the differences were emphasized the next day.

**Figure 4.**
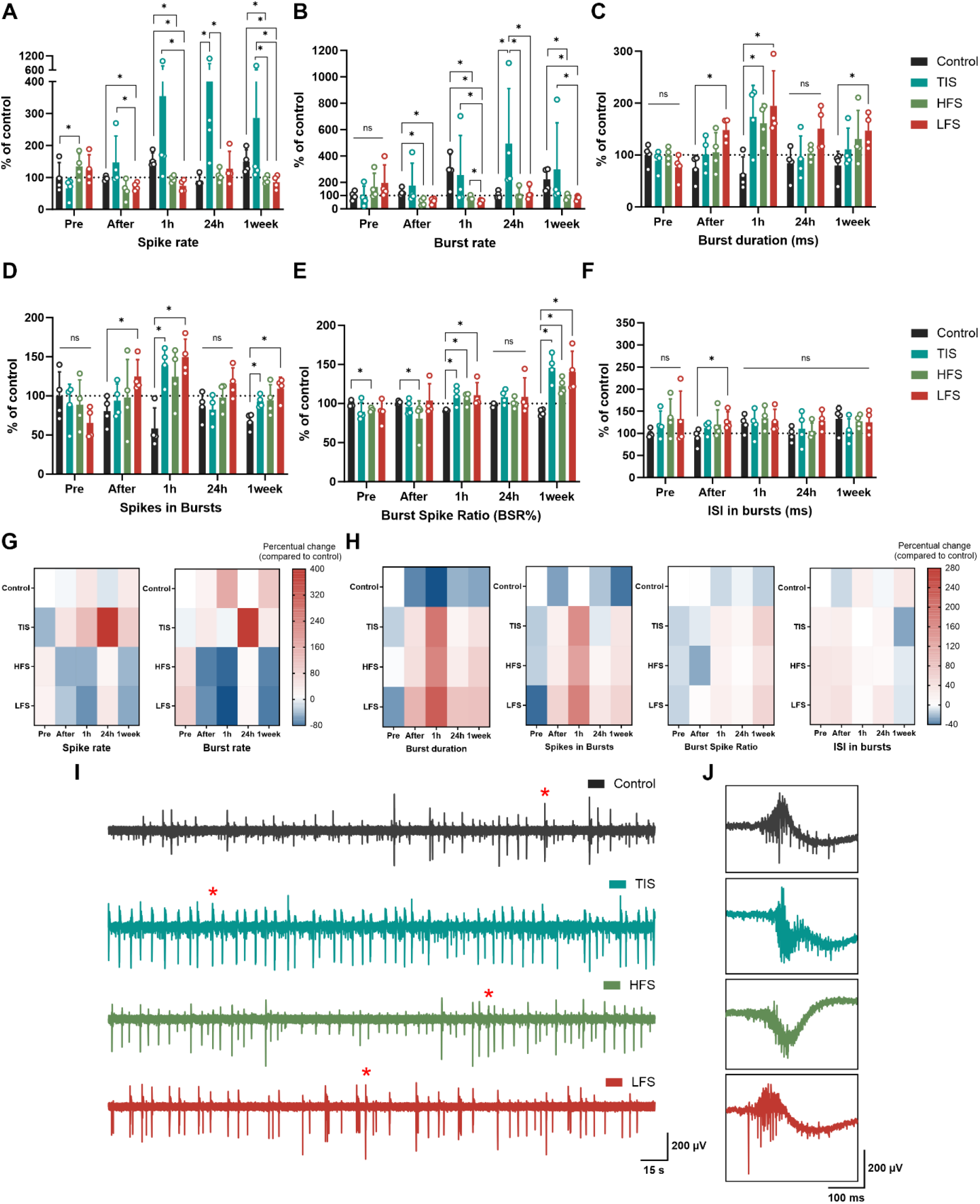
Electrophysiology of neuron cultures. (**A-F**) The effects of ES were compared to changes in control cultures at each time point. TIS (653+643 Hz with 10 Hz beat frequency) affected spike and burst rates by increasing the electrophysiological activity, especially after 24 hours. The effects of HFS (653 Hz) and LFS (10 Hz) were the most prominent one hour after the stimulation, as the spike and burst activities were suppressed and the duration of bursts prolonged. The burst spike ratio (BSR%) was increased for all ES types one hour after the stimulation, as were spikes in bursts for TIS and LFS. Some of the effects observed in spike rates and BSRs were observed even one week after the stimulation for TIS, HFS, and LFS. Bar graphs show means±SD with data points. (**G-H**) Heatmaps of the percentual increases and decreases compared to the control/sham cultures. Control values were adjusted to 0 (indicating no change) and values below that show decreases and values above 0 show increases in the respective activity parameters. (**I-J**) Representative signal traces of control, TI, HF, and LF-stimulated neurons 24 hours after the stimulation and their 300 ms cut-outs, indicated with red stars in the full signals. ns=not significant, ^*^p<0.05. (n=4 MEAs/condition. All statistical tests were done using the Mann-Whitney U test).

Twenty-four hours after the stimulation, TIS cultures had significantly increased spike rates and burst rates by +397 and +389%, respectively (**Figure 4A-B,G**; *p*=0.0286 for both). There were no differences in electrical activity for the LFS and HFS cultures after 24 hours compared to control cultures, but the differences in spike rates and burst rates were significant compared to TI-stimulated cultures (*p*=0.0286). Burst durations, spikes in bursts, BSRs, and ISI in bursts remained unchanged for all the stimulation conditions 24 hours after the stimulation (**Figure 4C-F,H**). Cultures were also assessed one week after the stimulation, and LFS and HFS cultures still showed decreased spike (-40 and -34%, respectively) and burst rates (-61 and - 57%, respectively) compared to control and TI-stimulated cultures (*p*=0.0286 for all). Also, burst duration (+86%) was higher for the LFS, as were the number of spikes in bursts and BSR (+60% and 58%, respectively, *p*=0.0286). Furthermore, TI-stimulated cultures showed elevated numbers of spikes in bursts and increased BSRs one week after the stimulation compared to control cultures (+36 and +64%, *p*=0.0286). The ISI in bursts remained unchanged for all one week after the stimulation (**Figure 4F**).

Taken together, TI-stimulation was the only form of ES applied that activated the neurons by increasing their spiking and bursting activity (**Figure 4G**). The temporally interfering electrical fields activated the neurons, and the trend was already visible one hour after the stimulation, but the effect was emphasized 24 hours after the stimulation. On the contrary, the “conventional” LFS suppressed neuronal electrical activity instead of eliciting it, and the effect was visible directly after stimulation and lasted up to a week (**Figure 4H**). Instead, the HFS had less of an impact on the neuronal cultures, but it seemed to impede electrical activity, especially one hour and one week after the applied stimulus. The percentual changes compared to the changes in the control cultures are depicted in **Figure 4G** for the spike rates and burst rates and in **Figure 4H** for the rest of the parameters.

The representative signal traces 24 hours after the stimulation for all the culture conditions are presented in **Figure 4I-J**. The TI-stimulated cultures appeared more electrically active than the other culture conditions, with repetitive spiking between the bursts. Moreover, the functional connectivity changes 24 hours after the stimulation, when the effects were the most prominent, were evaluated using CorSE analysis. **Supplementary Figure S2B** represents the CorSE figure of a representative MEA of each condition 24 hours after the stimulation and the results of the analysis. The connectivity results indicated that none of the stimulation methods affected the connectivity strengths of the cultures despite the changes in neuronal spiking and bursting, especially due to the TIS.

### Temporal interference stimulation of neuron-astrocyte co-cultures *in vitro*

After evaluating the effects of TIS, HFS, and LFS on neurons, we assessed the effects of TIS on neuron-astrocyte co-cultures that were stimulated, like neurons at DIV28 (n=4 MEAs/condition). All results and, the statistical tests used, and the details can be found in **Supplementary Tables S2-4**. Neurons and astrocytes were cultured at a ratio of approximately 1:1, and it was expected that astrocytes would modulate the neuronal electrophysiological response when subjected to TIS. At baseline (pre-stimulation), the cultures had very equal activities, and no differences were observed in any of the assessed parameters. Interestingly, astrocytes seemed to counteract the effects of TIS. In fact, there were no changes in the spike rates of the co-cultures in any of the recordings after the stimulation (**Figure 5A**). Furthermore, in contrast to neuronal cultures, the burst rate decreased 24 hours after the stimulation by 60% compared to the control cultures (**Figure 5B**, *p*=0.0286; Mann-Whiney U test). Interestingly, the burst duration was significantly prolonged for TI-stimulated cultures one week after the stimulation (+76%; **Figure 5C**, *p*=0.0286). In addition, the number of spikes in bursts increased one hour and one week after stimulation by 142% and 38%, respectively (**Figure 5D**; *p*=0.0286 for both). Also, the BSR% was elevated by 32%, and ISI in bursts decreased by -59% for the co-cultures following the TIS (**Figure 5E-F**; *p*=0.0286). However, compared to neuronal cultures without added astrocytes, the effect of TIS was relatively small, implying that astrocytes modulated the extracellular environment of neurons during and after TIS and, hence, counteracted the stimulation effects.

**Figure 5.**
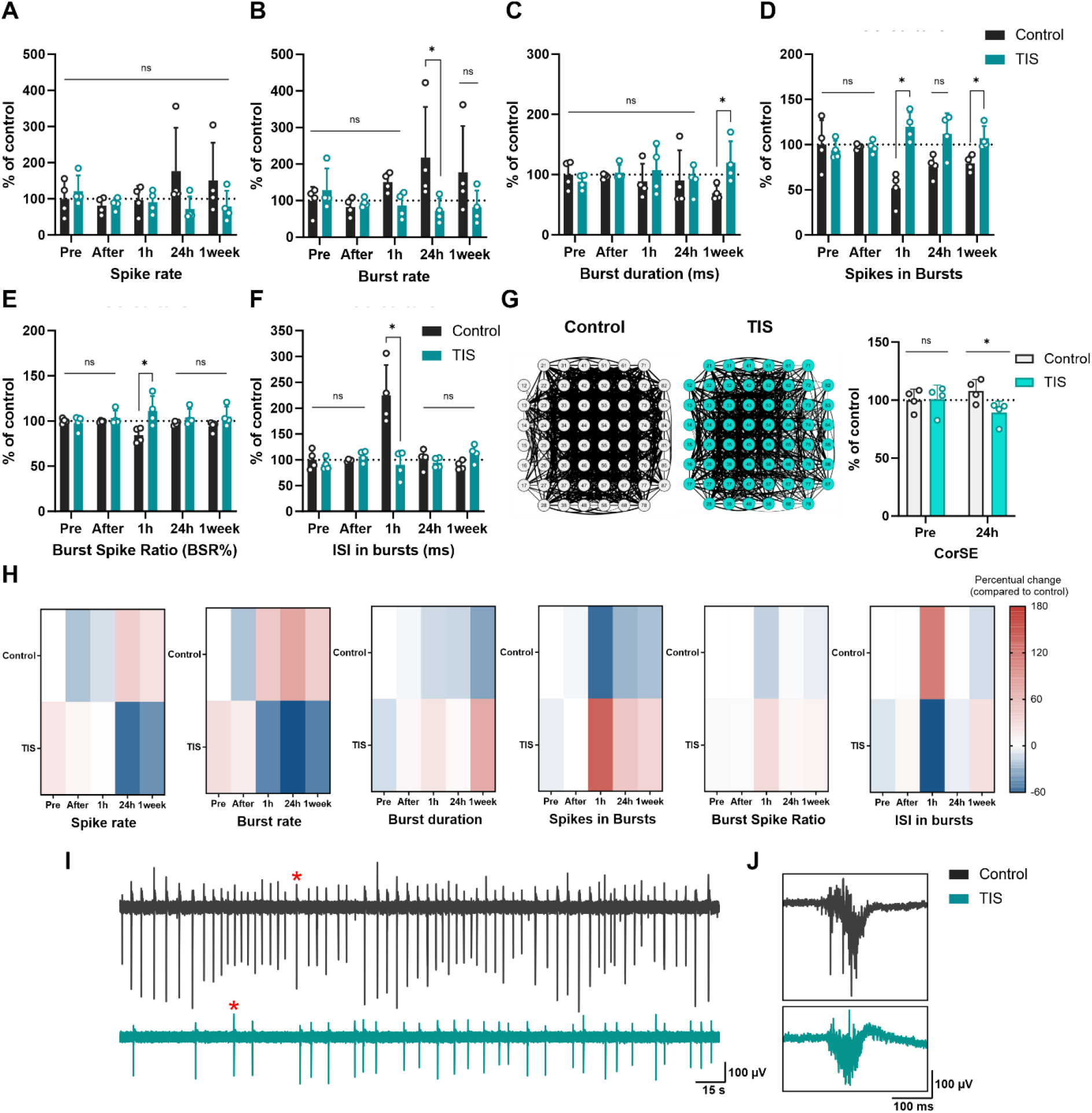
Electrophysiology of the co-cultures. (**A-F**) TIS did not affect the characterized electrical activity parameters in co-cultures, except for spikes in bursts and BSR% (increase) and burst duration (decrease) one hour after the stimulation. Also, burst rate decreased for the co-cultures 24 hours after the TIS compared to the control cultures. One week after the stimulation, burst duration and spikes in bursts were elevated for TI-stimulated cultures compared to control. (**G**) TIS decreased the connection strengths in the cultures 24 hours after the stimulation. (**H**) Heatmaps of the percentual increases and decreases compared to the control/sham cultures. Control values were adjusted to 0 (indicating no change); values below that show decreases, and values above 0 show increases in the respective activity parameters. (**I-J**) Representative activity signal traces of control and TI-stimulated co-cultures 24 hours after the stimulation. 300-ms cut-outs in (**J**) indicated with a red star in the full five-minute signals represented in (**I**). ns=not significant, ^*^p<0.05. (n=4 MEAs/condition. Bar graphs show means±SD with data points. All statistical tests were done using Mann-Whitney U test).

Next, we evaluated the effects of TIS on the functional connectivity of the co-cultures. The CorSE analysis was done for the co-cultures pre-stimulation and 24 hours after stimulation, when the effects of TIS were the most prominent for neurons. Intriguingly, TI-stimulated co-cultures experienced a slight decrease in connectivity compared to control cultures 24 hours after stimulation, as TIS reduced the connection strengths in the networks compared to the control by -17% (*p*=0.0286) (**Figure 5G**). Hence, although TIS had a relatively small effect on the electrophysiology of the co-cultures, the astrocytes and neurons seemed to respond by pruning the neuronal connection strengths formed in the networks. Representative signal traces of the control and TI-stimulated co-cultures 24 hours after the stimulation are presented in **Figure 5I-J**. The TI-stimulated co-cultures had much less spiking and bursting activity compared to the control cultures.

## Discussion

In this paper, we present a novel electrical field interference (EFI) stimulation system designed for *in vitro* neural cultures featuring temporally interfering electric fields. Temporal interference stimulation (TIS) utilizes high-frequency electric fields with slightly differing frequencies that superimpose spatially and temporally to create a low-frequency beating amplitude electric field. Since the concept was first introduced by Grossman et al., (2017) in the field of transcranial stimulation, it has sparked interest among neuroscience research communities. Previous TIS studies, aimed at optimizing parameters and gaining deeper insights into the underlying mechanisms of TIS, have been conducted primarily using animal or human models (Carmona-Barrón et al., 2023; Iszak et al., 2023; Ma et al., 2022; Song et al., 2021; Violante et al., 2023; Zhu et al., 2022) and computational simulations (Bahn et al., 2023; Lee et al., 2022a, 2022b, 2020; Mirzakhalili et al., 2020; Rampersad et al., 2019). In this study, we introduced an experimental *in vitro* setup that enabled us to explore the effects of TIS on neuronal cell electrophysiology at the cellular and network levels. TIS promises selective targeting of different spatial areas of neural populations, while at the same time minimizing off-target effects. The effects of TIS were compared to control cultures that were not subject to any stimulation and to other stimulation modalities that mimic the characteristics of TIS through distinct high- or low-frequency stimulation (HFS and LFS, respectively). The EFI setup used in this research was implemented by combining the DC-STIMULATOR MC (neuroConn GmbH, Ilmenau, Germany) and MEA2100 system (Multi Channel Systems GmbH, Reutlingen, Germany) with an in-house stimulus electrode ensemble.

We showed, with actual real-time recordings, that the resulting TIS signal exhibits 10 Hz amplitude envelopes and high-frequency components, as in (Grossman et al., 2017) *in vivo*. Notably, our results show that the TIS fields did not appear as a completely uniform signal across the electrode area. The local field strength derived from the TIS MEA recordings shows that the EFI field is strongest in the middle of the electrode area – as expected. The EFI field is the stimulation field sensed by neurons. Specifically, amongst the 60 electrodes and different field directions, the local electric potential differences were highest between two vertically neighboring electrodes and six horizontally neighboring electrodes in the center of the electrode array – a result fitting nicely with the theoretical consideration of the EFI. These findings indicate a selectivity of the introduced TIS system in targeting specific stimulation areas. Particularly with conventional stimulation methods, the local electric potential differences were the highest in proximity to the stimulation electrodes (Huang et al., 2017; Ineichen et al., 2018). Our results highlight the opportunity for TIS to produce a high local electrical field further from the stimulus electrodes the center. Hence, our results show that TIS can open new prospects for spatially targeted ES applications not only *in vivo* but also *in vitro*.

The biological effects of TIS were evaluated using two different culture types – neurons with minimum astrocyte support and co-cultures with equal proportions of neurons and astrocytes. We cultured the cells on microelectrode arrays (MEAs) and glass substrates and characterized the culture morphology, viability, and electrophysiological properties during maturation. The electrophysiological properties and other characteristics of the cultures, such as cell composition and morphology, resembled those in our studies using the same cell types (Ahtiainen et al., 2023, 2021). Immunofluorescent imaging showed that co-cultures exhibited a significantly higher proportion of astrocytes compared to neuronal cultures. Furthermore, both culture types displayed the presence of synapses, which is an indication of functional circuitry. Astrocytes enhanced the electrophysiological maturation of the cultures during the first weeks *in vitro*, but by DIV28, neuronal cultures were electrically more active, as the astrocytes seemed to suppress the repetitive firing of neurons, which we also observed in a previous study (Ahtiainen et al., 2021).

As hypothesized in the TIS stimulation theory, HFS at 653 Hz did not evoke increased electrophysiological responses in neurons, nor did the LFS at 10 Hz. Instead, HFS and LFS seemed to suppress neuronal spiking and bursting activity, and this suppressive effect seemed to persist even one week after stimulation. This suggests that LFS and HFS may have affected neurons, for example, by changing their excitability through long-term depression or potentiation and alterations in synaptic plasticity (Albensi et al., 2007). Furthermore, previous research suggests that the impact of HFS and LFS on neuronal activity is highly dependent on the specific stimulus site and parameters but that they could suppress epileptiform activity (Ahn et al., 2017; Lim et al., 2016; Paschen et al., 2020; Schiller and Bankirer, 2007). Moreover, emerging evidence suggests that high-frequency conduction blocks might block or inhibit the propagation of action potentials along axons, thereby contributing to the observed effects (Mirzakhalili et al., 2020). However, it is noteworthy to acknowledge that the ES parameters used, aspects of the experimental setup such as invasiveness, and consequently the results usually vary highly across studies, both *in vitro* and *in vivo*.

Our stimulation system effectively generated a TIS signal capable of modulating the neural cells cultured on top of the microelectrodes. The TIS signal comprised high-frequency components corresponding to HFS and a 10 Hz amplitude envelope similar to that produced by LFS. However, the spatial field strengths of LFS and TIS are far from equal. Thus, unlike LFS, the TIS elicited robust electrical responses in neurons, which were the most prominent 24 hours after stimulation. With computational methods, it has been demonstrated that the axon does not respond to the high-frequency components when the HF stimulus frequencies are the same (Δf=0), but it follows the envelope of the input signal in TIS (Karimi et al., 2019). In some cultures, the spike rates and burst rates increased over 1000% compared to control cultures, and they increased almost 400% on average. Varying neuronal responses to TIS were shown by Grossman et al., (2017) *in vivo* and Mirzakhalili et al., (2020) *in silico*, including activation or inactivation. Intriguingly, in our hands, while neuronal cultures appeared to activate in response to TIS, the co-cultures exhibited an opposing response. Therefore, based on our results, we could conclude that whether the response is active or inactive seems to depend also on the astrocyte impact in the cultures. Moreover, Mirzakhalili et al. (2020) proposed that the TI-activation of neurons via low-frequency envelopes requires an ion-channel-mediated signal rectification process and specific gating properties of the fast sodium (Na^+^) channels. Given the significant impact of astrocytes on the neuronal extracellular space and ion buffering, taken together with our results, these factors may contribute to the observed inactivation of neurons during TIS in neuron-astrocyte co-cultures. Another important intracellular messenger for astrocytes is calcium (Ca^2+^), which could regulate neuronal excitability in response to TIS by suppressing both (Agulhon et al., 2008; Bernardinelli et al., 2011; Ma et al., 2021). Therefore, for future experiments, Ca^2+^ imaging of astrocytes during TIS, HFS, and LFS could shed light on this matter.

While astrocytes seem to have notable implications for neuronal electrical activity in co-cultures during TIS, our findings suggest that astrocytes modulate the neuronal network during and following stimulation. The functional connectivity analysis (CorSE) indicated a reduction in connection strengths within co-cultures 24 hours after TIS. Additionally, our result aligns with a recent *in vivo* result found in humans. Violante et al. (2023) observed a similar reduction in functional connectivity after TIS with a beat frequency of 5 Hz (Violante et al., 2023). Therefore, neurons and astrocytes may have responded to TIS by rewiring their networks at synapses (Eroglu and Barres, 2010). The reasons for the observed impact in co-cultures are multifaceted – starting with the parameters of the stimulation, including the applied stimulation current and duration. Most brain stimulation studies, including the original study by Grossmann et al. (2017), use a 20-minute stimulation time, as it has been shown to induce direct effects and aftereffects in electroencephalograms (EEGs) (Kasten et al., 2016; Neuling et al., 2013). However, astrocytes may have caused the observed effects by counteracting the stimulation by regulating neuronal ion homeostasis.

In summary, we successfully established an *in vitro* setup for non-invasive stimulation of neuron and neuron-astrocyte co-cultures via temporally interfering electric fields. We demonstrated the capability of TIS to provide spatially targetable stimulation field *in vitro*. Interestingly, TIS with a precise spatial focus on the neuronal area of the MEA electrodes inflicted more pronounced effects in cultures with fewer astrocytes, resulting in a prominent electrophysiological response. TIS offers reduced side effects and, shown in this study, better spatial control over the stimulation, making it an attractive subject of study for future brain stimulation methods *in vitro*. This innovation holds promise for minimizing unwanted effects on non-targeted brain regions and assessing the interactions between neural cells and EFI stimulation for brain modulation purposes. Our study provides insight into the therapeutic potential of different stimulation systems and the effects of different stimulation paradigms on neural cells.

## Methods

### Cell culture

#### Microelectrode array and substrate preparations

MEAs were prepared similarly to (Ahtiainen et al., 2021). Briefly, MEAs (60MEA200/30iR-Tigr; Multi Channel Systems MCS GmbH, Germany) were coated with poly-d-lysine (0.1 mg/ml), rinsed with Dulbecco’s Phosphate Buffered Saline (DPBS), air dried, and subsequently coated with laminin (L2020, 20 µg/ml). MEAs were stored at +4 °C and taken to room temperature (RT) at least 30 minutes before cell plating. Each MEA consisted of 59 recording electrodes and one internal reference electrode in an 8x8 grid, each electrode having a 30 µm diameter and 200 µm electrode spacing. A coating protocol similar to that used for MEAs was applied for µwells (80807; ibidi GmbH, Gräfelfing, Germany) and glass coverslips for live/dead (L/D) and immunocytochemistry (ICC), respectively.

#### Cell plating on MEAs

Rat cortical neurons and astrocytes were plated onto MEAs according to a previously published protocol (Ahtiainen et al., 2023, 2021). Briefly, rat cortical astrocytes (N7745100, Thermo Fisher Scientific, Waltham, MA, USA) were treated to prevent proliferation with cytosine β-D-arabinofuranoside (ara-c; C1768, 2.5 µM, Sigma-Aldrich, St. Louis, MO, USA) and frozen in liquid nitrogen until plated in co-culture with neurons. On the day of plating, neurons (A1084001 or A1084002, Thermo Fisher Scientific) and arac-treated astrocytes were thawed, viability was determined with a Countess Automated Cell Counter (Thermo Fisher Scientific), and neurons and astrocytes were plated so that each MEA had 80,000 neurons and either no added astrocytes (later referred to as “neurons” for simplicity) or the same number of astrocytes and neurons (∼1:1), herein referred to as co-culture. In the same way, neurons or co-cultures were seeded into coverslips and µwells for further characterization, but the cell numbers were ½ or ¼ of what was used in MEAs, respectively. Neuron cultures were maintained in neurobasal plus medium with 2% B-12 Plus supplement, 1% P/S, and 1% GlutaMAX supplement, and the same medium was used for co-cultures in addition to 1% sodium pyruvate (all purchased from Thermo Fisher Scientific). Cells were fed by replacing at least half of the medium three times a week, and always after recording electrical activity. Cell growth and morphology were constantly monitored in the laboratory using a Nikon Eclipse Ts2 (Nikon Corporation, Japan) microscope.

#### Immunocytochemistry

ICC was performed to characterize the cultures at DIV29. The full ICC protocol is available in (Ahtiainen et al., 2021). The primary antibodies used were glial fibrillary acidic protein (GFAP; AB5804, rabbit, 1:1000), microtubule associated protein 2 (MAP2; PA1-10005, chicken, 1:1000 or 1:2000), postsynaptic density protein 95 (PSD-95; MA1-045, mouse, 1:200), and synaptophysin (SYN; MA5-14532, rabbit, 1:50). The secondary antibodies used were goat anti-mouse 488 (A32723; 1:500), goat anti-Rabbit 555 (A-21428; 1:500), and goat anti-chicken 647 (A32933; 1:500). All the antibodies were purchased from Thermo Fisher Scientific (Waltham, MA, USA), except for AB5804, which was purchased from Sigma-Aldrich (St. Louis, MO, USA), and all were diluted in 5% (v/v) goat serum (Sigma–Aldrich) in Dulbecco’s Phosphate-Buffered Saline (DPBS). Samples were stained with DAPI (4′,6-diamidino-2-phenylindole; D1306, Thermo Fisher Scientific) prior to mounting (10-15 min at RT). Samples were mounted with ProLong™ Gold Antifade Mountant (P10144, Thermo Fisher Scientific; 24 hours in the dark at RT). The coverslips were stored at +4°C until imaging with an Olympus IX51 fluorescence microscope with an Olympus DP30BW camera (Olympus Corporation, Hamburg, Germany). Fiji (ImageJ, National Institute of Health, USA) software was used to process the images.

#### Live/dead assay

A live/dead viability/cytotoxicity kit (L3224; Thermo Fisher Scientific) with ethidium homodimer-1 (4 µM) and calcein-AM (2 µM) was used according to the manufacturer’s instructions for both cultures in µwells at DIV28. The images were taken using an Olympus IX51 fluorescence microscope and analyzed using a particle size–based analysis similar to that of (Ahtiainen et al., 2021).

### Stimulation setup and system validation

MEAs were connected to the DC-STIMULATOR MC (neuroConn GmbH, Germany), which provided the stimulation currents to MEAs via platinum electrodes. MEA signals from neurons were recorded prior to, during, and after stimulation with the MEA2100 system (Multi Channel Systems MCS GmbH), which enabled real-time monitoring of the effects of stimulus on neurons. The stimulus was carried through a special 3D-printed cap composed of Biomed Amber Resin (Formlabs GmbH, Germany). The cap was printed according to the MEA dimensions to establish a suitable electric field using a stereolithography 3D printer Form 3B+ (Formlabs GmbH, Germany). The cap was designed using Solidworks 2021 (Dassault Systèmes SolidWorks Corp., France), and various cap designs were tested and validated prior to the final design. The cap consisted of four elevated holes on the top for the application of platinum rod electrodes submerged in the cell medium during stimulation. Additionally, the cap had two air holes for a ventilation tube (0.3 mbar; 5% CO_2_+19% O_2_+75% N_2_). The design of the cap is represented in **Supplementary Figure S1A-C**. Holes for the platinum rods (1.2x0.8 mm) at a 25° angle for the appropriate TIS conductance were designed, and for the stimulation, four bar-shaped platinum electrodes (1 mmx0.5 mmx12 mm) were connected with 1.5 mm diameter connectors (DIN 42802-2) that were soldered to the platinum electrodes. The generated currents for TIS and high frequencies were initially confirmed with measurements using an oscilloscope (MP720665 from Multicomp Pro, Premier Farnell Ltd, USA) (**Supplementary Figure S1D-E**), as well as thereafter with the MEA2100 system, which was also used during the experiments to record electrical activity.

For the TIS, frequencies of 653 Hz and 643 Hz were chosen for stimulation, as they are the highest possible prime numbers below 700 Hz (3 dB cut-off frequency of stimulator) with a 10 Hz difference between them. Prior to the experiment, different stimulation paradigms were evaluated. With a 250 µA current, different stimulation times (1 min, 5 min, and 20 min) were tested, and five minutes was agreed to be an appropriate stimulation time (with a 30 second fade in/fade out time for the stimulation itself). Next, we tested different stimulation currents of 250 µA, 350 µA, and 450 µA for five minutes of stimulation. As the highest tested stimulation current of 450 µA did not affect cell viability, it was used in the final experiments.

### Stimulation and MEA recordings

#### Experimental setup

Neurons and neuron-astrocyte co-cultures (∼1:1) were cultured on MEAs for 28 days *in vitro* (DIV). At DIV28, neurons were stimulated with TIS, HFS, and LFS or didn’t receive any stimulation (control/sham). MEAs were always allowed to settle in the preamplifier for five minutes before the first recording. For the stimulation, MEAs were not moved from the recording site until the first stimulation was conducted directly after recording the baseline electrical activity. ES (or sham) was applied for five minutes. For those MEAs that did not receive any stimulation, the stimulation was simulated by keeping the MEAs outside the incubator in the MEA headstage for the same amount of time as those MEAs that were stimulated (TIS/HFS/LFS). After the ES (or sham), at least half of the cell medium was replaced, and MEAs were put back in the incubator. Electrical activity was recorded for five minutes before sham/stimulation and immediately after, one hour after, 24 hours after, and one week after ES (or sham). Data were collected and analyzed by comparing the stimulation effects to the percentual change in the control cultures at each time point in question. Co-cultures of neurons and astrocytes were either stimulated with TIS or were not stimulated (control/sham), and analyzed similarly. Each ES condition per culture had 4 MEAs (a total of 24 MEAs for neurons and co-cultures).

TIS was achieved by combining two stimulation frequencies (f_1_=643 Hz, f_2_=653 Hz) with 10 Hz differences to create 10 Hz amplitude envelopes corresponding to the frequency of LFS. The purpose of using LFS and HFS for the neurons was to compare the stimulation paradigms alone to enable evaluation of the effects of interference stimulation with respect to control cultures without stimulation. Detailed stimulation paradigms are available in **Table 3**.

**Table 3.**
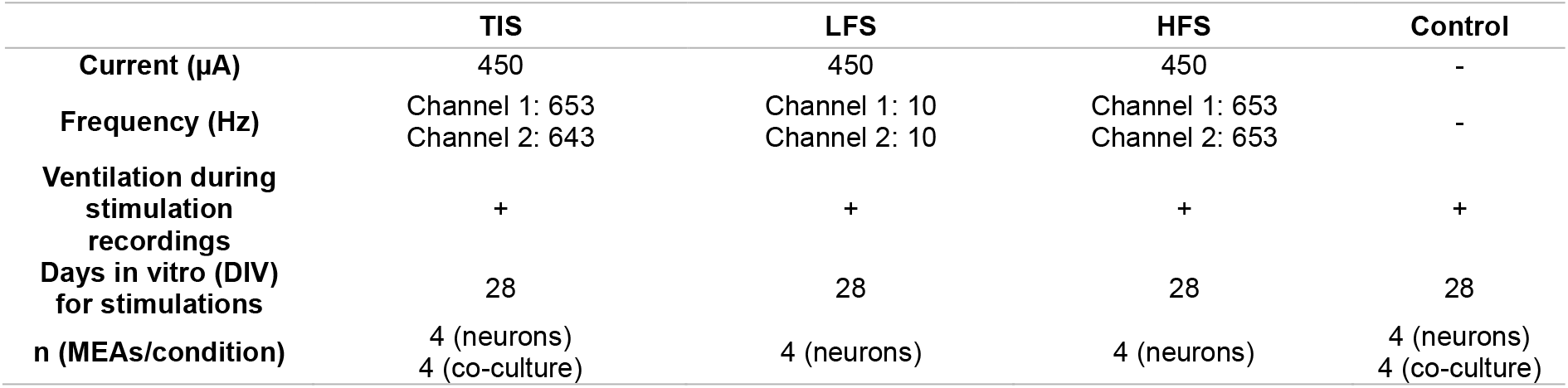
Stimulation paradigms in the study.

### Data analysis

#### MEA data

MEA electrophysiological data were recorded with a sampling rate of 50 kHz using the MEA2100-System and Multichannel Experimenter software (both from Multichannel Systems MCS GmbH, Reutlingen, Germany).

Raw electrophysiological data were further analyzed with Matlab 2022a (MathWorks, Inc., Natick, MA, USA) using a previously published, widely used tool for spike sorting (Chaure et al., 2018), also implemented in (Ahtiainen et al., 2021). Briefly, a second-order bandpass (300–3000 Hz) elliptic filter with a threshold of ±5σ (σ=median (|*χ*|/0.6745), x=bandpass filtered signal) and 1.5 second detector dead time after spike was used for sorting. From the sorted, filtered signals, we analyzed the timestamps of the signals and other parameters of the electrical activity, such as spike rate, burst rate, burst duration, spikes in bursts, burst spike ratio (BSR%), and interspike interval (ISI) in bursts (details in **Table 2**). These values were extracted from the data using a tool developed previously in our group (Kapucu et al., 2012; Välkki et al., 2017). Signal traces were extracted from the analyzed data and plotted with Matlab 2022a.

To observe stimulation electric field strengths, bipolar signals were calculated from the filtered MEA data recorded during the stimulation of co-culture MEA for the TI-stimulation. Bipolar signals were calculated by subtracting the signals from two adjacent microelectrodes for all microelectrodes (c.f., **Figure 2B** for the MEA layout noting the 90° rotation) in both horizontal and vertical directions, resulting in two sets of bipolar signals. The bipolar signals were converted to field strengths, given the microelectrode distance of 200 µm. The two sets of TIS field strengths and the microelectrodes are illustrated in **Figure 2F**. For all the stimulation modalities (TIS/HFS/LFS/control), the signals and field strengths were calculated by using the same methods for unfiltered raw data derived from neuronal cultures during stimulation/sham (**Supplementary Figure S3**).

#### CorSE Analysis

Correlated spectral entropy (CorSE) analysis was used to evaluate the connectivity strength of the networks. CorSE uses correlations of time-varying spectral entropies between all MEA channels to evaluate synchronicity. The connectivity strength between MEA electrodes is established by their magnitude of correlation, and the overall average connection strength considers all the electrode pairs in the network. The analysis was done according to (Kapucu et al., 2016), using Matlab 2022a with a connection threshold of 0.75. CorSE was analyzed for both neurons and co-cultures at DIV28 and DIV29, 24 hours after the stimulation/sham.

### Statistical analysis

All statistical analysis was done with Graphpad Prism (v.9.0). Significance values below 0.05 were considered significant. Asterisks in the figure indicate the level of significance, where ^*^*p*<0.05, ^**^*p*<0.01, ^***^*p*<0.001, and *ns* are not significant. All electrophysiological data were statistically tested with the Mann-Whitney U test due to the non-normal distribution of the data. For the effects of ES (% of control), the results were obtained by comparing the changes in each group with the respective changes in the control group at the time of the recording in question. Hence, the percentual changes were obtained by comparing the changes caused by the stimulation to the respective changes in the control cultures occurring at each time point. All statistical details, as well as absolute values for the electrical data, are available in the Supplementary Material (**Supplementary Tables S1–S4**).

## Data and code availability

The data can be made available upon request by the corresponding author.

## Acknowledgements

The authors acknowledge the doctoral school of Faculty of Medicine and Health Technology, Tampere University, for supporting the work of A.A. and the Tampere Imaging Facility for their services. The authors acknowledge the Academy of Finland -funded Centre of Excellence in Body-On-Chip Research (grant number 353178) for supporting laboratory equipment. Otherwise, this research did not receive any specific grant from funding agencies in the public, commercial, or not-for-profit sectors.

## Author contributions

A.A., J.A.K.H., J.H., and A.H., conceived and designed the study. A.A. and L.L. conducted the experiments; A.A. plated the cells and L.L. helped to maintain them in the laboratory. A.A. performed cell assays and immunocytochemistry. A.A. and L.L. conducted the stimulations. A.A. and L.L. designed the final cap design, and L.L. drew and printed it. A.A. analyzed and displayed the results. J.M.A.T. generated the original MATLAB code for electric potential differences to calculate the V/m. The study was directed and coordinated by J.A.K.H, J.H, and A.H. A.A. compiled and wrote the paper with input from all authors.

