## Supplemental Information for "Stimulation of Neurons and Astrocytes via Temporally Interfering Electric Fields"

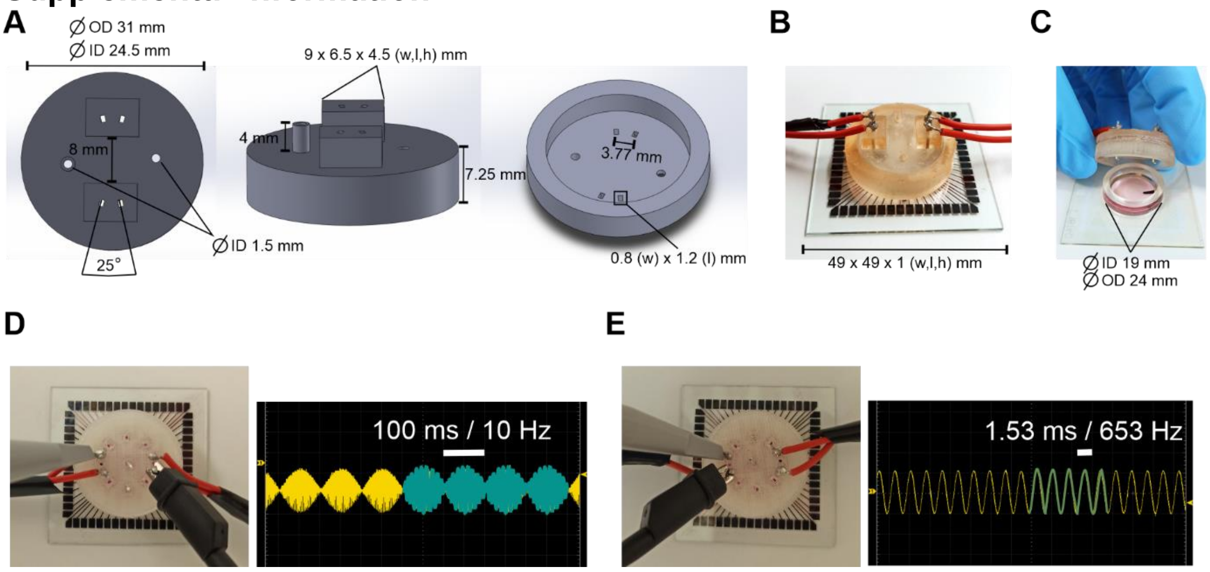


***Supplementary Figure S1.*** *(**A**) The top, side, and under-views of the 3D-printed cap. The cap had elevated holes for the gas tube and the platinum electrode pairs, as well as a small hole for gas exchange. (**B**) A MEA with a cap and electrodes in place. (**C**) The platinum electrodes were submerged in the cell medium at even distances and depths. (**D**) Oscilloscope measurements of the TIS signal at 500 µA. TIS envelopes are indicated in blue. (**E**) Oscilloscope measurements of high-frequency signals at 500 µA. High-frequency signals are indicated in green.*

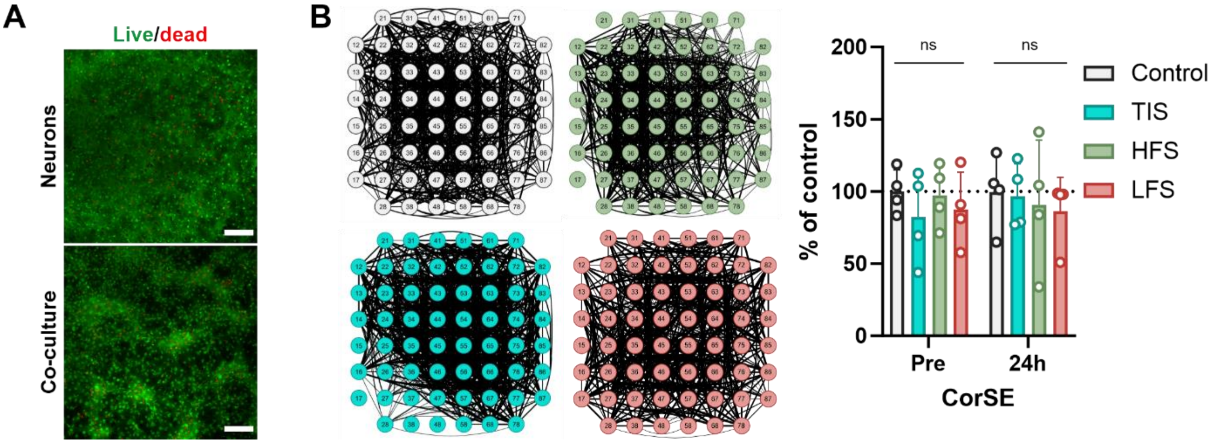


***Supplementary Figure S2.*** *(**A**) Live/dead assay of neuron and neuron-astrocyte co-cultures at DIV28. There were no differences in the viability of the cultures. The scale bar is 200 µm. (**B**) CorSE analysis 24 hours after the stimulation. There were no differences in the connectivity between the groups. ns=not significant (n=4 MEAs/condition. Mann-Whitney U test was used for the CorSE analysis).*

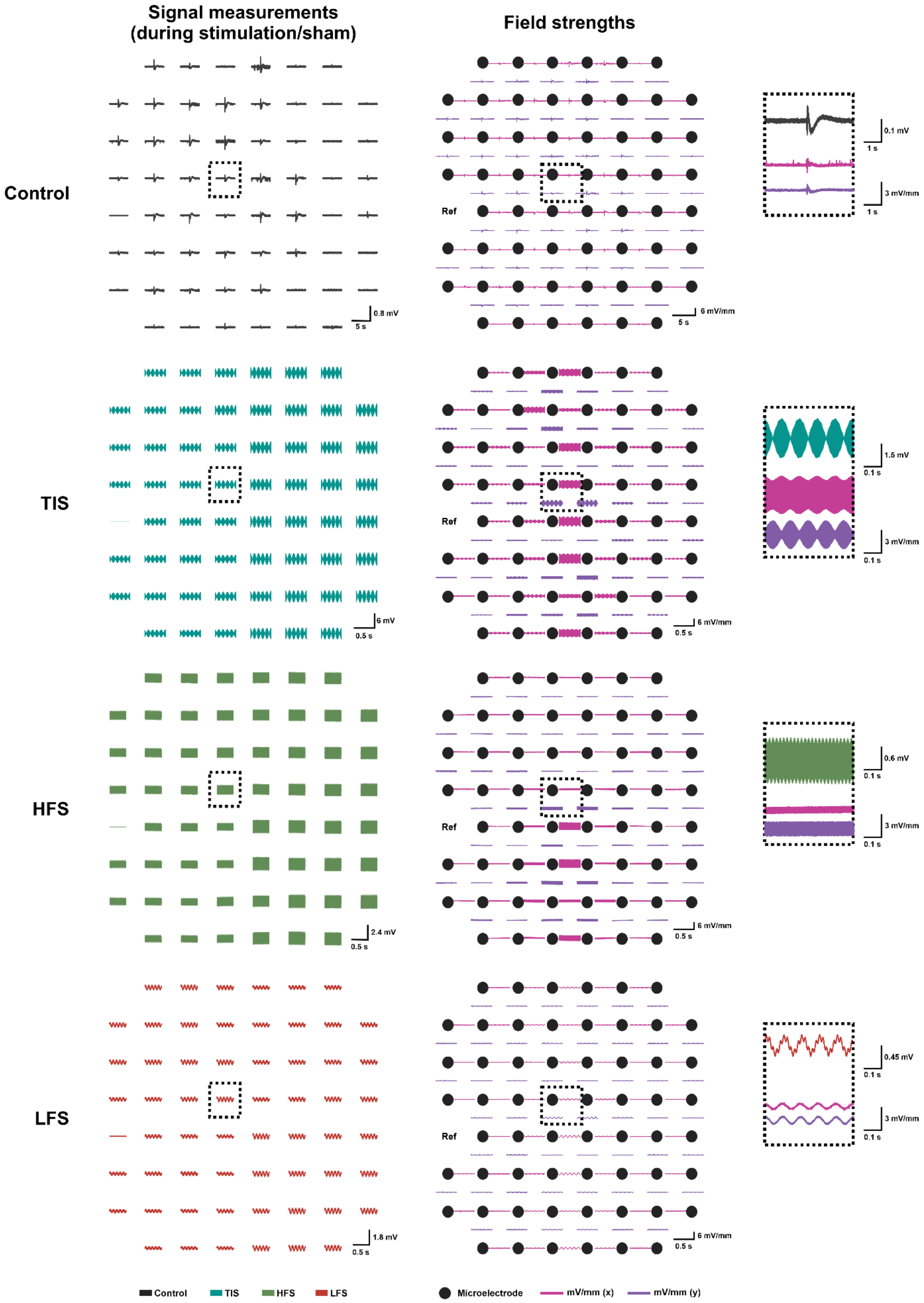


***Supplementary Figure S3. Representative signal measurements and field strengths for control, TIS, HFS and LFS.*** *TI-stimulated cultures experienced the highest field strengths (mV/mm) especially in the middle of the electrode area. Instead, HFS and LFS cultures had much lower and uniform field strengths. Close-ups for one electrode are visible in the right side. (All signals are derived from unfiltered data from neuronal cultures).*

***Supplementary Table S1.*** *Maturation of the cultures. Mann-Whitney U test, comparison between neurons and co-cultures.*

### Electrophysiology – maturation (%)

**Mann-Whitney-U-test**

**Significance (median1, median2, U), *p***

**Neurons vs. co-culture**

| | Spike rate | Burst rate | Burst duration | Spikes In Bursts | Burst Spike Ratio | ISI in bursts | Active electrodes |
|---|---|---|---|---|---|---|---|
| **DIV7** | (80.60, 118.2, 26), **0.0192 *** | (2.640, 4.038, 26), **0.0192 *** | (180.1, 290.2, 28), **0.0274 *** | (18.97, 17.35, 51), 0.4523 | (0.3349, 0.504, 6), **<0.0001 ***** | (20.65, 17.66, 47), 0.3196 | (55, 59, 28), **0.0158 *** |
| **DIV14** | (410.2, 342.7, 42), 0.1924 | (9.025, 4.958, 30), **0.0382 *** | (175.0, 185.6, 48), 0.3503 | (33.80, 65.24, 2), **<0.0001 ***** | (0.7252, 0.9598, 0), **<0.0001 ***** | (7.194, 3.238, 1), **<0.0001 ***** | (59, 59, 64), >0.9999 |
| **DIV21** | (722.2, 614.4, 57), 0.6968 | (14.74, 15.22, 53), 0.5283 | (142.7, 100.3, 27), **0.0230 *** | (41.35, 36.81, 49), 0.3826 | (0.8682, 0.9416, 6), **<0.0001 ***** | (4.040, 2.981, 20), **0.0057 **** | (59, 59, 64), >0.9999 |
| **DIV28** | (760.5, 411.8, 16), **0.0022 **** | (17.03, 10.96, 28), **0.0274 *** | (143.5, 100.2, 21), **0.0071 **** | (33.54, 34.27, 59), 0.7875 | (0.7513, 0.9447, 4), **<0.0001 ***** | (5.398, 3.073, 2), **<0.0001 ***** | (59, 59, 60), >0.9999 |
| **DIV35** | (1112, 563.7, 0), **0.0286 *** | (28.34, 16.62, 0), **0.0286 *** | (127.8, 78.57, 3), 0.2000 | (24.87, 29.96, 6), 0.6857 | (0.7237, 0.9157, 0), **0.0286 *** | (6.337, 3.031, 0), **0.0286 *** | (59, 59, 7.5), >0.9999 |

### Electrophysiology - maturation

**Mean ± SD**

**Neurons**

| | Spike rate (spikes/min) | Burst rate (bursts/min) | Burst duration (ms) | Spikes In Bursts | Burst Spike Ratio (0-1) | ISI in bursts (ms) | Active electrodes (0-60 (icl.ref)) |
|---|---|---|---|---|---|---|---|
| **DIV7** | 92±75 | 3±1 | 184±119 | 20±22 | 0.3±0.1 | 25±16 | 46±17 |
| **DIV14** | 453±209 | 12±9 | 164±43 | 34±11 | 0.7±0.1 | 8±4 | 59±0 |
| **DIV21** | 751±273 | 16±6 | 136±23 | 41±9 | 0.8±0.1 | 4±2 | 59±0 |
| **DIV28** | 860±371 | 21±15 | 137±31 | 34±11 | 0.8±0.1 | 6±2 | 59±0 |
| **DIV35** | 1144±350 | 32±13 | 119±46 | 27±11 | 0.7±0.0 | 6±2 | 55±8 |

**Co-culture**

| | | | | | | | |
|---|---|---|---|---|---|---|---|
| **DIV7** | 144±74 | 4±1 | 258±87 | 22±11 | 0.5±0.1 | 19±8 | 59±0 |
| **DIV14** | 352±60 | 6±1 | 181±27 | 65±12 | 1.0±0.0 | 3±1 | 59±0 |
| **DIV21** | 646±124 | 16±4 | 110±26 | 39±7 | 0.9±0.0 | 3±1 | 59±0 |
| **DIV28** | 428±172 | 12±5 | 101±20 | 35±7 | 0.9±0.1 | 3±1 | 59±0 |
| **DIV35** | 490±195 | 15±4 | 73±13 | 28±6 | 0.9±0.1 | 3±0.0 | 56±6 |

***Supplementary Table S2.*** *Stimulation results. Mann-Whitney U test, comparisons done by comparing the stimulation results to control change at each time point in question.*

| **Electrophysiology – stimulation (%)**<br>**Mann-Whitney-U-test** | | | | | |
|---|---|---|---|---|---|
| **Significance (median1, median2, U), *p*** | | | | | |
| **Spike rate** | | | | | |
| **Neurons** | **Pre** | **After** | **1h** | **1d** | **1w** |
| **ctrl vs. TIS** | (86.48, 77.69, 5), 0.4857 | (96.67, 111.8, 4), 0.3429 | (145.6, 168.8, 6), 0.6857 | (81.75, 263.8, 0), **0.0286 *** | (141.3, 128.3, 7), 0.8857 |
| **ctrl vs. HFS** | (86.48, 131.3, 4), 0.3429 | (96.67, 61.57, 2), 0.1143 | (145.6, 95.46, 0), **0.0286 *** | (81.75, 103.0, 3), 0.200 | (141.3, 91.16, 0), **0.0286 *** |
| **ctrl vs. LFS** | (86.48, 114.3, 4), 0.3429 | (96.67, 71.37, 0), **0.0286 *** | (145.6, 71.90 6), **0.0286 *** | (81.75, 112.3, 6), 0.6857 | (141.3, 86.94, 0), **0.0286 *** |
| **TIS vs. HFS** | (77.69, 131.3, 0), **0.0286 *** | (111.8, 61.57, 1), 0.0571 | (168.8, 95.46, 1), 0.0571 | (263.8, 103.0, 0), **0.0286 *** | (128.3, 91.16, 0), **0.0286 *** |
| **TIS vs. LFS** | (77.69, 114.3, 2), 0.1143 | (111.8, 71.37, 0), **0.0286 *** | (168.8, 71.90, 0), **0.0286 *** | (263.8, 112.3, 1), 0.0571 | (128.3, 86.94, 0), **0.0286 *** |
| **HFS vs. LFS** | (131.3, 114.3, 7), 0.8857 | (61.57, 71.37, 6), 0.6857 | (95.46, 71.90, 3), 0.200 | (103.0, 112.3, 6), 0.6857 | (91.16, 86.94, 6), 0.6857 |
| **Co-cultures** | | | | | |
| **ctrl vs. TIS** | (98.99, 106.6, 6), 0.6857 | (83.57, 87.74, 7), 0.8857 | (105.0, 91.89, 7), 0.8857 | (118.8, 55.83, 3), 0.2000 | (113.7, 67.37, 4), 0.3429 |
| **Burst rate** | | | | | |
| **Neurons** | | | | | |
| **ctrl vs. TIS** | (100.5, 73.48, 6), 0.6857 | (117.8, 101.9, 5), 0.4857 | (293.9, 118.5, 5), 0.4857 | (98.49, 326.8, 0), **0.0286 *** | (226.1, 132.4, 5), 0.4857 |
| **ctrl vs. HFS** | (100.5, 127.7, 4), 0.3429 | (117.8, 48.25, 0), **0.0286 *** | (293.9, 74.06, 0), **0.0286 *** | (98.49, 92.85, 6), 0.6857 | (226.1, 90.87, 0), **0.0286 *** |
| **ctrl vs. LFS** | (100.5, 141.1, 3), 0.2000 | (117.8, 66.45, 0), **0.0286 *** | (293.9, 52.42, 0), **0.0286 *** | (98.49, 104.6, 8), >0.9999 | (226.1, 86.79, 0), **0.0286 *** |
| **TIS vs. HFS** | (73.48, 127.7, 3), 0.2000 | (101.9, 48.25, 1), 0.0571 | (118.5, 74.06, 2), 0.1143 | (326.8, 92.85 0), **0.0286 *** | (132.4, 90.87, 1), 0.0571 |
| **TIS vs. LFS** | (73.48, 141.1, 3), 0.2000 | (101.9, 66.45, 1), 0.0571 | (118.5, 52.42, 0), **0.0286 *** | (326.8, 104.6 0), **0.0286 *** | (132.4, 86.79, 0), **0.0286 *** |
| **HFS vs. LFS** | (127.7, 141.1, 6), 0.6857 | (48.25, 66.45, 7), 0.8857 | (74.06, 52.42, 0), **0.0286 *** | (92.85, 104.6, 6), 0.6857 | (90.87, 86.79, 7), 0.8857 |
| **Co-cultures** | | | | | |
| **ctrl vs. TIS** | (113.1, 107.4, 8), >0.9999 | (85.15, 91.72, 6), 0.6857 | (153.0, 88.20, 1), 0.0571 | (162.0, 65.21 0), **0.0286 *** | (136.4, 70.21, 3), 0.2000 |
| **Burst duration** | | | | | |
| **Neurons** | | | | | |
| **ctrl vs. TIS** | (104.2, 95.54, 4), 0.3429 | (81.93, 101.0, 5), 0.4857 | (54.09, 191.6, 1), 0.0571 | (88.93, 83.21, 7), 0.8857 | (91.05, 94.74, 5), 0.4857 |
| **ctrl vs. HFS** | (104.2, 99.34, 7), 0.8857 | (81.93, 102.7, 5), 0.4857 | (54.09, 168.6, 0), **0.0286 *** | (88.93, 103.3, 5), 0.4857 | (91.05, 114.8, 3), 0.2000 |
| **ctrl vs. LFS** | (104.2, 81.31, 3), 0.2000 | (81.93, 151.3, 0), **0.0286 *** | (54.09, 165.9, 0), **0.0286 *** | (88.93, 146.9, 2), 0.1143 | (91.05, 146.2, 0), **0.0286 *** |
| **TIS vs. HFS** | (95.54, 99.34, 7), 0.8857 | (101.0, 102.7, 8), >0.9999 | (191.6, 168.6 6), 0.6857 | (83.21, 103.3, 5), 0.4857 | (94.74, 114.8, 6), 0.6857 |
| **TIS vs. LFS** | (95.54, 81.31, 5), 0.4857 | (101.0, 151.3, 2), 0.1143 | (191.6, 165.9 8), >0.9999 | (83.21, 146.9, 2), 0.1143 | (94.74, 146.2, 3), 0.2000 |
| **HFS vs. LFS** | (99.34, 81.31, 3), 0.2000 | (102.7, 151.3, 2), 0.1143 | (168.6, 165.9 6), 0.6857 | (103.3, 146.9, 2), 0.1143 | (114.8, 146.2, 6), 0.6857 |
| **Co-cultures** | | | | | |
| **ctrl vs. TIS** | (103.4, 90.52, 6), 0.6857 | (98.09, 96.79, 8), >0.9999 | (80.15, 104.9, 6), 0.6857 | (69.80, 93.88, 7), 0.8857 | (65.54, 109.5, 0), **0.0286 *** |
| **Spikes in Burst** | | | | | |
| **Neurons** | | | | | |
| **ctrl vs. TIS** | (94.57, 96.83, 7), 0.8857 | (82.07, 90.65, 5), 0.4857 | (48.97, 144.3, 0), **0.0286 *** | (89.00, 84.21, 6), 0.6857 | (70.37, 91.48, 0), **0.0286 *** |
| **ctrl vs. HFS** | (94.57, 85.90, 6), 0.6857 | (82.07, 96.25, 6), 0.6857 | (48.97, 131.2, 1), 0.0571 | (89.00, 101.8, 5), 0.4857 | (70.37, 90.97, 1), 0.0571 |

| | | | | | |
|---|---|---|---|---|---|
| **ctrl vs. LFS** | (94.57, 64.10, 3), 0.2000 | (82.07, 116.0, 0), **0.0286 *** | (48.97, 144.9, 0), **0.0286 *** | (89.00, 113.2, 1), 0.0571 | (70.37, 114.7, 0), **0.0286 *** |
| **TIS vs. HFS** | (96.83, 85.90, 8), >0.9999 | (90.65, 96.25, 7), 0.8857 | (144.3, 131.2, 6), 0.6857 | (84.21, 101.8, 4), 0.3429 | (91.48, 90.97, 7), 0.8857 |
| **TIS vs. LFS** | (96.83, 64.10, 3), 0.2000 | (90.65, 116.0, 3), 0.2000 | (144.3, 144.9, 7), 0.8857 | (84.21, 113.2, 1), 0.0571 | (91.48, 114.7, 3), 0.2000 |
| **HFS vs. LFS** | (85.90, 64.10, 4), 0.3429 | (90.65, 116.0, 4), 0.3429 | (144.9, 131.2 5), 0.4857 | (101.8, 113.2, 4), 0.3429 | (90.97, 114.7, 5), 0.4857 |
| **Co-cultures** | | | | | |
| **ctrl vs. TIS** | (100.7, 90.45, 6), 0.6857 | (98.11, 97.10, 8), >0.9999 | (56.36, 118.3, 0), **0.0286 *** | (78.22, 118.2, 1), 0.0571 | (79.38, 101.6, 0), **0.0286 *** |
| | | | **Burst Spike Ratio** | | |
| **Neurons** | | | | | |
| **ctrl vs. TIS** | (99.30, 85.89, 4), 0.3429 | (100.7, 94.29, 6), 0.6857 | (90.77, 112.3, 0), **0.0286 *** | (97.26, 108.6, 2), 0.1143 | (89.23, 147.2, 0), **0.0286 *** |
| **ctrl vs. HFS** | (99.30, 91.94, 0), **0.0286 *** | (100.7, 91.20, 0), **0.0286 *** | (90.77, 109.0, 0), **0.0286 *** | (97.26, 99.78, 6), 0.6857 | (89.23, 121.3, 0), **0.0286 *** |
| **ctrl vs. LFS** | (99.30, 90.29, 4), 0.3429 | (100.7, 94.92, 4), 0.3429 | (90.77, 103.9, 0), **0.0286 *** | (97.26, 99.48, 6), 0.6857 | (89.23, 132.3, 0), **0.0286 *** |
| **TIS vs. HFS** | (85.89, 91.94 4), 0.3429 | (94.29, 91.20, 6), 0.6857 | (112.3, 109.0, 6), 0.6857 | (108.6, 99.78, 4), 0.3429 | (147.2, 121.3, 2), 0.1143 |
| **TIS vs. LFS** | (85.89, 90.29, 7), 0.8857 | (94.29, 94.92, 6), 0.6857 | (112.3, 103.9, 7), 0.8857 | (108.6, 99.48, 5), 0.4857 | (147.2, 132.3, 5), 0.4857 |
| **HFS vs. LFS** | (91.94, 90.29, 6), 0.6857 | (91.20, 94.92, 4), 0.3429 | (109.0, 103.9, 7), 0.8857 | (99.78, 99.48, 8), >0.9999 | (121.3, 132.3, 4), 0.3429 |
| **Co-cultures** | | | | | |
| **ctrl vs. TIS** | (100.1, 100.5, 8), >0.9999 | (99.47, 99.90, 8), >0.9999 | (84.92, 108.2, 0), **0.0286 *** | (97.85, 100.8, 3), 0.2000 | (96.18, 101.8, 3), 0.2000 |
| | | | **ISI in bursts** | | |
| **Neurons** | | | | | |
| **ctrl vs. TIS** | (97.50, 115.9, 4), 0.3429 | (94.10, 116.1, 2), 0.1143 | (123.5, 124.1, 8), >0.9999 | (97.95, 113.6, 6), 0.6857 | (142.4, 101.6, 4), 0.3429 |
| **ctrl vs. HFS** | (97.50, 130.1, 4), 0.3429 | (94.10, 109.1, 3), 0.2000 | (123.5, 133.3, 6), 0.6857 | (97.95, 99.20, 7), 0.8857 | (142.4, 127.8, 5), 0.4857 |
| **ctrl vs. LFS** | (97.50, 109.4, 7), 0.8857 | (94.10, 121.7, 0), **0.0286 *** | (123.5, 118.6, 8), >0.9999 | (97.95, 127.8, 2), 0.1143 | (142.4, 126.6, 7), 0.8857 |
| **TIS vs. HFS** | (115.9, 130.1, 5), 0.4857 | (116.1, 109.1, 8), >0.9999 | (124.1, 133.3, 6), 0.6857 | (113.6, 99.20, 8), >0.9999 | (101.6, 127.8, 3), 0.2000 |
| **TIS vs. LFS** | (115.9, 109.4, 7), 0.8857 | (116.1, 121.7, 4), 0.3429 | (124.1, 118.6, 8), >0.9999 | (113.6, 127.8, 5), 0.4857 | (101.6, 126.6, 4), 0.3429 |
| **HFS vs. LFS** | (130.1, 109.4, 7), 0.8857 | (109.1, 121.7, 5), 0.4857 | (133.3, 118.6, 7), 0.8857 | (99.20, 127.8, 2), 0.1143 | (127.8, 126.6, 8), >0.9999 |
| **Co-cultures** | | | | | |
| **ctrl vs. TIS** | (98.42, 87.34, 7), 0.8857 | (99.01, 108.8, 4), 0.3429 | (209.0, 95.66, 0), **0.0286 *** | (105.3, 94.10, 4), 0.3429 | (87.45, 114.3, 2), 0.1143 |
| | | | **CorSE** | | |
| **Neurons** | | | | | |
| **ctrl vs. TIS** | (98.90, 86.49, 5), 0.4857 | | | (103.3, 93.53, 8), >0.9999 | |
| **ctrl vs. HFS** | (98.90, 99.13, 8), >0.9999 | | | (103.3, 94.04, 7), 0.8857 | |
| **ctrl vs. LFS** | (98.90, 86.04, 5), 0.4857 | | | (103.3, 97.79, 3), 0.2000 | |
| **TIS vs. HFS** | (86.49, 99.13 5), 0.4857 | | | (93.53, 94.04, 8), >0.9999 | |
| **TIS vs. LFS** | (86.49, 86.04, 7), 0.8857 | | | (93.53, 97.79, 6), 0.6857 | |
| **HFS vs. LFS** | (99.13, 86.04, 7), 0.8857 | | | (94.04, 97.79, 7), 0.8857 | |
| **Co-cultures** | | | | | |
| **ctrl vs. TIS** | (102.2, 105.3, 7), 0.8857 | | | (109.1, 93.39, 0), **0.0286 *** | |

***Supplementary Table S3.*** *Absolute values (mean±sd, standard error of mean [SEM]) of the electrical stimulation results.*

#### Electrophysiology - stimulation

| Mean ±SD, SEM | | | | | | |
|---|---|---|---|---|---|---|
| **Spike rate (spikes/min)** | | | | | | |
| | | **Pre** | **After** | **1h** | **1d** | **1w** |
| **Neurons** | Control | 809.2±376.2, 188.1 | 777.5±351.2, 175.6 | 1215±476.6, 238.3 | 713.6±290.7, 145.3 | 1144±349.9, 174.9 |
| | TIS | 531.6±249.8, 124.9 | 608.9±169.1, 84.53 | 1682±579.2, 289.6 | 1353±330.9, 165.5 | 1295±361.2, 180.6 |
| | HFS | 1086±320.2, 160.1 | 588.3±234.8, 117.4 | 1568±590, 295 | 1016±327.9, 164 | 1418±427.3, 213.7 |
| | LFS | 1012±370, 185 | 675.6±190.8, 95.41 | 1183±698.5, 349.3 | 1255±1021, 510.7 | 1258±714.4, 357.2 |
| **Co-cultures** | Control | 386.1±190.9, 95.44 | 306.1±159.6, 79.82 | 344.6±195.3, 97.64 | 553.4±148.6, 74.31 | 489.6±194.7, 97.35 |
| | TIS | 469.4±166.6, 83.32 | 324.4±138.2, 69.10 | 354±76, 38 | 448.2±140.4, 70.19 | 418.6±127.5, 63.73 |
| **Burst rate (bursts/min)** | | | | | | |
| **Neurons** | Control | 15.41±5.075, 2.537 | 19.31±6.595, 3.298 | 41.57±16.74, 8.369 | 15.58±3.517, 1.759 | 32.47±12.91, 6.456 |
| | TIS | 14.42±12.21, 6.105 | 19.27±7.762, 3.881 | 51.92±21.46, 10.73 | 44.65±17.55, 8.774 | 47.59±19.84, 9.919 |
| | HFS | 25.70±15.76, 7.880 | 14.08±5.515, 2.758 | 54.36±29.38, 14.69 | 26.92±12.35, 6.176 | 45.33±18.52, 9.260 |
| | LFS | 30.21±21.03, 10.51 | 18.88±4.970, 2.485 | 46.73±40.24, 20.12 | 43.42±49.41, 24.71 | 53.68±41.30, 20.65 |
| **Co-cultures** | Control | 10.19±3.829, 1.914 | 8.089±3.472, 1.736 | 15.16±6.333, 3.167 | 18.37±3.151, 1.575 | 14.83±4.411, 2.205 |
| | TIS | 13.12±6.054, 3.027 | 9.850±4.74, 2.37 | 15.02±3.245, 1.623 | 14.95±2.896, 1.448 | 13.28±3.684, 1.842 |
| **Burst duration (ms)** | | | | | | |
| **Neurons** | Control | 150.2±30.21, 15.10 | 108.7±34.10, 17.05 | 89.43±23.85, 11.92 | 123.2±42.90, 21.45 | 118.7±45.68, 22.84 |
| | TIS | 134.1±30.55, 15.28 | 95.15±29.53, 14.77 | 134.2±45.43, 22.72 | 96.75±24.97, 12.49 | 110.7±10.46, 5.230 |
| | HFS | 149.2±21.90, 10.95 | 109.7±27.05, 13.52 | 139.8±18.22, 9.112 | 125.9±25.19, 12.59 | 147.9±40.38, 20.19 |
| | LFS | 114.7±36.42, 18.21 | 121.9±43.48, 21.74 | 123.2±19.95, 9.973 | 138.3±53.95, 26.98 | 130.3±47.69, 23.85 |
| **Co-culture** | Control | 107.5±25.08, 12.54 | 103.9±21.08, 10.54 | 92.98±27.94, 13.97 | 92.82±42.47, 21.23 | 73.38±13.12, 6.560 |
| | TIS | 94.87±13.28, 6.64 | 93.57±7.710, 3.855 | 84.81±20.07, 10.03 | 72.75±13.53, 6.763 | 75.85±14.54, 7.270 |
| **Spikes in Burst** | | | | | | |
| **Neurons** | Control | 40.06±12.33, 6.163 | 31.12±7.804, 3.902 | 21.61±4.831, 2.415 | 34.59±11.95, 5.976 | 27.39±10.95, 5.474 |
| | TIS | 35.05±10.96, 5.482 | 24.61±6.907, 3.454 | 25.88±7.607, 3.803 | 24.23±7.066, 3.533 | 22.52±8.025, 4.012 |
| | HFS | 35.48±12.75, 6.377 | 24.57±8.971, 4.485 | 21.99±2.633, 1.317 | 29.08±7.983, 3.992 | 21.68±3.667, 1.833 |
| | LFS | 26.27±7.814, 3.907 | 24.92±5.921, 2.961 | 20.63±3.961, 1.980 | 25.59±5.090, 2.545 | 19.27±5.111, 2.556 |

| | | | | | | |
|---|---|---|---|---|---|---|
| **Co-culture** | Control | 36.42±9.891, 4.945 | 35.47±8.883, 4.441 | 18.01±6.131, 3.065 | 27.12±5.240, 2.62 | 28.34±5.598, 2.799 |
| | TIS | 34.22±3.847, 1.924 | 32.25±2.078, 1.039 | 20.05±1.162, 0.5811 | 28.25±5.348, 2.674 | 28.38±3.546, 1.773 |
| **Burst Spike Ratio (0-1)** | | | | | | |
| **Neurons** | Control | 0.8171±0.0303, 0.0151 | 0.8254±0.0357, 0.0179 | 0.7431±0.0251, 0.0126 | 0.7957±0.0236, 0.0118 | 0.7248±0.0332, 0.0169 |
| | TIS | 0.7339±0.0979, 0.0489 | 0.7043±0.0710, 0.0355 | 0.7304±0.0322, 0.0161 | 0.7687±0.0541, 0.0271 | 0.7290±0.1115, 0.0557 |
| | HFS | 0.7519±0.0194, 0.0097 | 0.6129±0.1737, 0.0868 | 0.7279±0.0570, 0.0285 | 0.7374±0.0678, 0.0339 | 0.6299±0.0487, 0.0243 |
| | LFS | 0.7314±0.117, 0.0585 | 0.7532±0.0858, 0.0429 | 0.7252±0.0484, 0.0242 | 0.7573±0.0985, 0.0492 | 0.6937±0.1121, 0.0561 |
| **Co-culture** | Control | 0.9441±0.0274, 0.0137 | 0.9399±0.0315, 0.0158 | 0.7972±0.0848, 0.0424 | 0.9207±0.0285, 0.0143 | 0.8889±0.0726, 0.0363 |
| | TIS | 0.9229±0.075, 0.0375 | 0.9458±0.0154, 0.0077 | 0.8604±0.0827, 0.0413 | 0.9319±0.0152, 0.0076 | 0.9053±0.0303, 0.0151 |
| **ISI in bursts (ms)** | | | | | | |
| **Neurons** | Control | 4.741±0.4361, 0.218 | 4.333±1.218, 0.6088 | 5.778±1.091, 0.5454 | 4.626±0.8505, 0.4253 | 6.323±1.713, 0.8565 |
| | TIS | 5.689±1.448, 0.7239 | 5.888±1.805, 0.9026 | 8.592±3.219, 1.610 | 6.371±3.553, 1.777 | 8.307±4.258, 2.129 |
| | HFS | 6.399±2.037, 1.018 | 6.905±2.175, 1.087 | 9.996±1.549, 0.7743 | 6.409±1.603, 0.8016 | 10.73±3.064, 1.532 |
| | LFS | 6.284±2.990, 1.495 | 7.097±2.610, 1.305 | 9.351±3.276, 1.638 | 8.224±5.239, 2.619 | 10.30±4.370, 2.185 |
| **Co-culture** | Control | 3.401±0.6343, 0.3172 | 3.359±0.6093, 0.3047 | 7.410±0.8927, 0.4463 | 3.419±0.6958, 0.3479 | 2.991±0.3460, 0.1730 |
| | TIS | 3.104±0.3923, 0.1962 | 3.229±0.0738, 0.0369 | 5.978±1.514, 0.7568 | 2.921±0.1997, 0.0998 | 3.023±0.1510, 0.0755 |
| **CorSE (0-1)** | | | | | | |
| **Neurons** | Control | 0.68±0.10, 0.05 | | | 0.67±0.18, 0.09 | |
| | TIS | 0.56±0.22, 0.11 | | | 0.52±0.19, 0.10 | |
| | HFS | 0.66±0.15, 0.07 | | | 0.62±0.31, 0.16 | |
| | LFS | 0.60±0.18, 0.09 | | | 0.54±0.25, 0.12 | |
| **Co-culture** | Control | 0.78±0.07, 0.04 | | | 0.84±0.05, 0.02 | |
| | TIS | 0.78±0.10, 0.05 | | | 0.75±0.09, 0.05 | |

***Supplementary Table S4.*** *Other results (viability and CorSE).*

| Other results<br>Mann-Whitney-U-test | | |
|---|---|---|
| Significance (median1, median2, U), *p* | | |
| Neurons vs. co-culture | | |
| | **Viability DIV28** | **CorSE DIV28** |
| **Neurons vs. co-culture** | (1.032, 1.015, 84), 0.3766 | (0.6289, 0.8139, 20), **0.0057 **** |
| **Mean±SD** | | |
| | **Viability DIV28** | **CorSE DIV28** |
| **Neurons** | 83±10 | 0.62±0.16 |
| **Co-culture** | 85±2 | 0.78±0.08 |
